# MYCBP interacts with Sakura and Otu and is essential for germline stem cell renewal and differentiation and oogenesis

**DOI:** 10.1101/2025.07.01.662550

**Authors:** Azali Azlan, Ryuya Fukunaga

**Affiliations:** Department of Biological Chemistry, Johns Hopkins School of Medicine, Baltimore, MD 21205

## Abstract

The self-renewal and differentiation of germline stem cells (GSCs) are tightly regulated during oogenesis. The *Drosophila* female germline provides a powerful model to study these regulatory mechanisms. We previously identified Sakura (also known as Bourbon/CG14545) as a crucial factor for maintenance and differentiation of GSCs and oogenesis, and demonstrated that Sakura binds to Ovarian Tumor (Otu), another essential regulator of these processes. Here, we identify MYCBP (c-Myc binding protein) as an additional essential component of this regulatory network. We show that MYCBP physically associates with itself, Sakura, and Otu, forming binary and ternary complexes including a MYCBP•Sakura•Otu complex. MYCBP is highly expressed in the ovary, and *mycbp* null mutant females exhibit rudimentary ovaries with germline-less and tumorous ovarioles, fail to produce eggs, and are completely sterile. Germline-specific depletion of *mycbp* disrupts Dpp/BMP signaling, causing aberrant expression of *bag-of-marbles* (*bam*) and leading to defective differentiation and GSC loss. In addition, *mycbp* is required for female-specific splicing of *sex-lethal* (*sxl*), a master regulator of sex identity determination. These phenotypes closely resemble those observed those of *sakura* and *otu* mutants. Together, our findings reveal that MYCBP functions in concert with Sakura and Otu to coordinate self-renewal and differentiation of GSCs and oogenesis in *Drosophila*.

## Introduction

Oogenesis—the process by which germline stem cells (GSCs) develop into mature female gametes (oocytes)—is governed by multiple layers of regulation involving numerous genes. GSCs maintain their undifferentiated state while undergoing asymmetric division to produce one self-renewing GSC and one differentiating daughter cell, known as a cystoblast. Disruption of this balance can lead to either GSC loss, impairing oocyte production, or the overproliferation of undifferentiated cells, resulting in tumorous phenotypes and compromised fertility (Gateff et al. 1996; Lin 1997; Cox et al. 1998; Ohlstein et al. 2000). While precise control of GSC self-renewal and cystoblast differentiation is essential for proper oogenesis, the underlying molecular mechanisms remain incompletely understood, and additional regulatory factors likely remain to be identified.

The fruit fly *Drosophila melanogaster*, a genetically tractable model organism, has long served as a powerful system for studying GSC regulation and and oogenesis research (Kirilly and Xie 2007). *Drosophila* females possess a pair of ovaries, each composed of 12-16 ovarioles. At the anterior tip of each ovariole is a structure known as the germarium, which typically houses two to three GSCs (Fig 1A). These GSCs reside at the anterior-most region, in direct contact with cap cells and escort cells that form a specialized niche required for GSC maintenance (Lin and Spradling 1993; de Cuevas and Matunis 2011; Losick et al. 2011). Upon asymmetric mitotic division, a GSC produces a daughter GSC and a cystoblast, which subsequently undergoes four rounds of mitotic divisions with incomplete cytokinesis to form a 16-cell cyst. Of these 16 interconnected cells, one becomes the oocyte, and the remaining 15 differentiate into polyploid nurse cells that supply the oocyte with essential RNAs and proteins.

**Fig 1.**
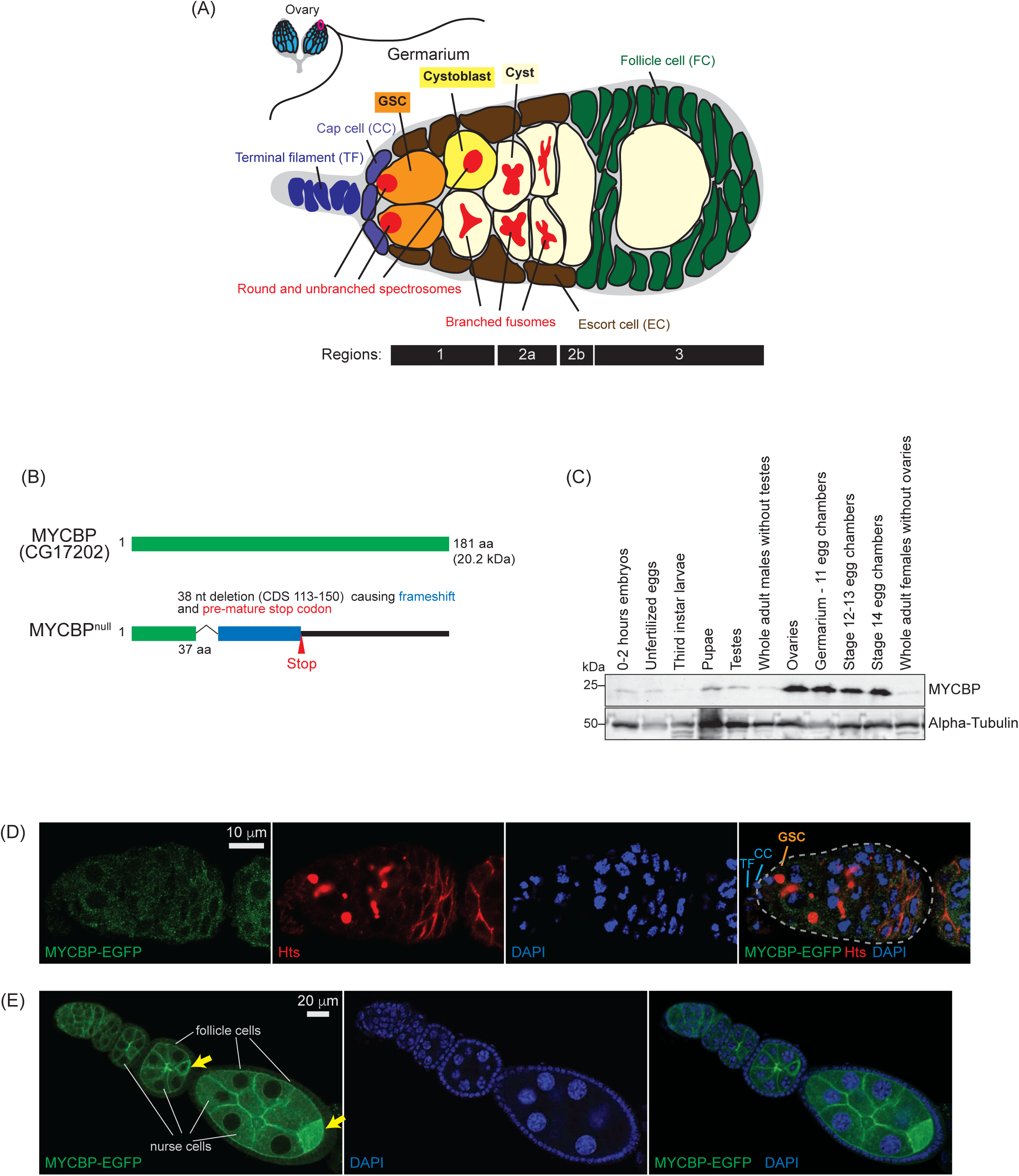
MYCBP expression pattern and mutant allele. (A) Schematic illustration of *Drosophila* ovary and germarium. Each female has a pair of ovaries, each consisting of 12-16 ovarioles (cyan). The germarium (outlined in magenta) is located at the anterior tip of each ovariole and contains both germ cells and somatic cells. Germ cells include germline stem cells (GSCs), cystoblasts, cysts, and differentiating oocytes. Somatic cells include terminal filament (TF) cells, cap cells (CCc), escort cells (ECs), and follicle cells (FCs). GSCs and cystoblasts have spherical, unbranched spectrosomes, whereas cysts posesss branched fusomes. The distinct regions of the germarium—1, 2a, 2b, and 3—are indicated. (B) *Drosophila* MYCBP (CG17202) protein and the null mutant allele generated in this study. (C) Western blot of dissected fly tissues. (D) Confocal images of germaria from *mycbp-EGFP* transgenic flies. MYCBP-EGFP (green), Hts (red), and DAPI (blue). Scale bar: 10 μm. (E) Confocal images of egg chambers from *mycbp-EGFP* transgenic flies. MYCBP-EGFP (green) and DAPI (blue). MYCBP-EGFP is highly expressed in nurse cells, enriched in the developing oocyte (yellow arrow), and is also detectable in somatic follicle cells. Scale bar: 20 μm.

We recently identified a novel gene, *sakura* (also known as *bourbon/CG14545*) as an essential factor for GSC self-renewal and differentiation and oogenesis in *Drosophila* (Azlan et al. 2024). The Sakura protein, consisting of 114 amino acids, is exclusively expressed in female germline cells, including GSCs. We found that Sakura physically interacts with Ovarian Tumor (Otu), a known regulator of these same processes. Mutations in either *sakura* or *otu* result in tumorous germline overgrowth, germ cell loss, defects in oocyte specification, and aberrations in sexual identity determination, including the failure of female-specific splicing of *sex-lethal* (*sxl*) mRNA and the consequent production of the male-specific isoform (Smith and King 1966; Gans et al. 1975; Gollin and King 1981; King and Riley 1982; Storto and King 1988; Pauli et al. 1993; Xu and Rubin 1993; Rodesch et al. 1997; Glenn and Searles 2001; Azlan et al. 2024). Otu is known to form a deubiquitinase complex with Bag-of-marbles (Bam), a key differentiation factor, and to deubiquitinate Cyclin A (CycA), thereby, stabilizing CycA and promoting GSC differentiation (Ji et al. 2017). In addition, Otu has been shown to bind RNA (Ji et al. 2019). However, the detailed molecular mechanisms by which Otu and Sakura function in GSC self-renewal and differentiation and oogenesis remain poorly understood.

In this study, we identify c-Myc binding protein (MYCBP, CG17202) (Fig 1B), an uncharacterized 181-amino-acid protein and the *Drosophila* ortholog of human c-Myc-binding protein, as a critical factor for GSC self-renewal and differentiation and oogenesis. We show that MYCBP physically associates with itself, Sakura, and Otu, forming binary and ternary complexes including a MYCBP•Sakura•Otu complex. MYCBP is highly expressed in female germline cells and is required for GSC self-renewal and differentiation, oogenesis, and female-specific splicing of *sxl* mRNA. Strikingly, *mycbp*, *sakura*, and *otu* mutants exhibit similar phenotypes. Our findings suggest that MYCBP functions together with Sakura and Otu as a critical module governing GSC regulation and oogenesis.

## Results

### MYCBP is highly expressed in the ovaries

MYCBP was one of the five proteins identified by mass spectrometry following co-immunoprecipitation with Sakura-EGFP from ovary lysates, suggesting a physical interaction between Sakura and MYCBP (Table 1) (Azlan et al. 2024). To examine MYCBP protein expression, we generated polyclonal anti-MYCBP antibodies against the recombinant full-length MYCBP protein. Western blot using this antibody revealed that MYCBP protein is highly expressed in the ovary, including germarium-stage 11 egg chambers, stage 12-13 egg chambers and stage 14 egg chambers (Fig 1C).

**Table 1.** Number of unique peptide counts detected by mass-spec of Sakura-EGFP co-IP samples. Proteins with at least two unique peptide signals present in all three biological replicates of the Sakura-EGFP samples with no unique peptide signal in any of the three biological replicates of the negative control or only one unique peptide signal in only one of the three biological replicates of the negative control are shown.

| Identified protein name | Unique peptide counts |  |  |  |  |  |
| --- | --- | --- | --- | --- | --- | --- |
|  | Sakura-EGFP samples |  |  | w1118 samples<br>(negative control) |  |  |
|  | Replicate<br>1 | Replicate<br>2 | Replicate<br>3 | Replicate<br>1 | Replicate<br>2 | Replicate<br>3 |
| Ovarian tumor (Otu) | 44 | 18 | 38 | 0 | 0 | 0 |
| CG4679 | 17 | 2 | 8 | 0 | 0 | 0 |
| CG14997 | 17 | 2 | 7 | 0 | 0 | 0 |
| mRpS22 | 11 | 2 | 6 | 0 | 1 | 0 |
| MYCBP | 4 | 2 | 4 | 0 | 1 | 0 |

### MYCBP is highly expressed in germ cells and cytoplasmic

To analyze MYCBP expression and localization, we created transgenic flies expressing a MYCBP-EGFP fusion protein under the control of the *mycbp* promoter. Oogenesis begins in the germarium, which contains 2-3 GSCs, identifiable by round, unbranched spectrosomes in contact with cap cells (Fig 1A) (Kirilly and Xie 2007). In contrast, developing cysts exhibit branched fusomes. We used hu-li tai shao (HTS) antibody to visualize both spectrosomes and fusomes (Fig 1D). Confocal imaging showed that MYCBP*-*EGFP localizes to the cytoplasm of germ cells, including GSCs, cysts, nurse cells, and developing oocytes (Fig 1D and 1E). Within egg chambers, MYCBP-EGFP was expressed highly in germline cells (nurse cells and developing oocytes) and was enriched in developing oocytes, while it was also detectably expressed in somatic follicle cells (Fig 1E).

### MYCBP forms complexes with Otu and Sakura

To identify MYCBP-interacting proteins, we performed co-immunoprecipitation using anti-GFP beads on ovary lysates from MYCBP-EGFP-expressing flies, followed by mass spectrometry. Ovary lysates from *w1118* flies served as negative controls. Mass spectrometry identified Otu as the top interactor and also detected Sakura (Table 2).

**Table 2.** Abundance of Proteins identified by mass-spec of MYCBP-EGFP co-IP samples. Proteins with adjusted p-value <0.05 or those detected in all MCYBP-EGFP replicates but not in any w1118 replicated are shown.

| Identified<br>protein<br>name | Abundance |  |  |  |  |  | log2(Fold-Change) | Adjusted<br>p-value |
| --- | --- | --- | --- | --- | --- | --- | --- | --- |
|  | MYCBP-EGFP samples |  |  | w1118 samples (Negative<br>controls) |  |  |  |  |
|  | Replicate<br>1 | Replicate<br>2 | Replicate<br>3 | Replicate<br>1 | Replicate<br>2 | Replicate<br>3 |  |  |
| MYCBP | 29.93 | 29.53 | 30.54 | 16.69 | nd | 21.67 | 10.82 | 1.4E-11 |
| Otu | 26.16 | 25.28 | 25.68 | nd | 18.57 | 20.74 | 6.05 | 0.009 |
| Sakura | 19.25 | 18.68 | 17.27 | nd | nd | nd | NA | NA |
| Sec71 | 26.94 | 26.28 | 27.22 | nd | nd | nd | NA | NA |
| CG15715 | 24.42 | 22.41 | 22.58 | nd | nd | nd | NA | NA |
| Rg | 22.61 | 22.31 | 23.16 | nd | nd | nd | NA | NA |
| mRpS2 | 21.30 | 19.51 | 20.08 | nd | nd | nd | NA | NA |
| CG7878 | 20.33 | 17.92 | 17.86 | nd | nd | nd | NA | NA |
| Elav | 19.98 | 18.26 | 19.10 | nd | nd | nd | NA | NA |
| EIF4G2 | 19.86 | 18.45 | 18.94 | nd | nd | nd | NA | NA |
| Rbp9 | 19.60 | 18.16 | 18.86 | nd | nd | nd | NA | NA |
| Whd | 19.58 | 17.81 | 18.50 | nd | nd | nd | NA | NA |
| Cht10 | 19.45 | 18.36 | 18.85 | nd | nd | nd | NA | NA |
| CG31717 | 18.82 | 17.39 | 18.39 | nd | nd | nd | NA | NA |
| RhoGAP15B | 18.72 | 18.40 | 19.29 | nd | nd | nd | NA | NA |
| Spg7 | 17.98 | 16.94 | 17.35 | nd | nd | nd | NA | NA |
| Polr3C | 17.79 | 23.48 | 21.61 | nd | nd | nd | NA | NA |
| Adck1 | 16.38 | 14.35 | 15.44 | nd | nd | nd | NA | NA |
| CG4332 | 15.74 | 13.69 | 13.82 | nd | nd | nd | NA | NA |

Western blotting confirmed that both Otu and Sakura co-immunoprecipitate with MYCBP-EGFP in ovary lysates (Fig 2A). Additionally, endogenous MYCBP co-immunoprecipitated with endogenous Otu and Sakura from wild-type ovary lysates (Fig 2B), confirming these interactions in vivo.

**Fig 2.**
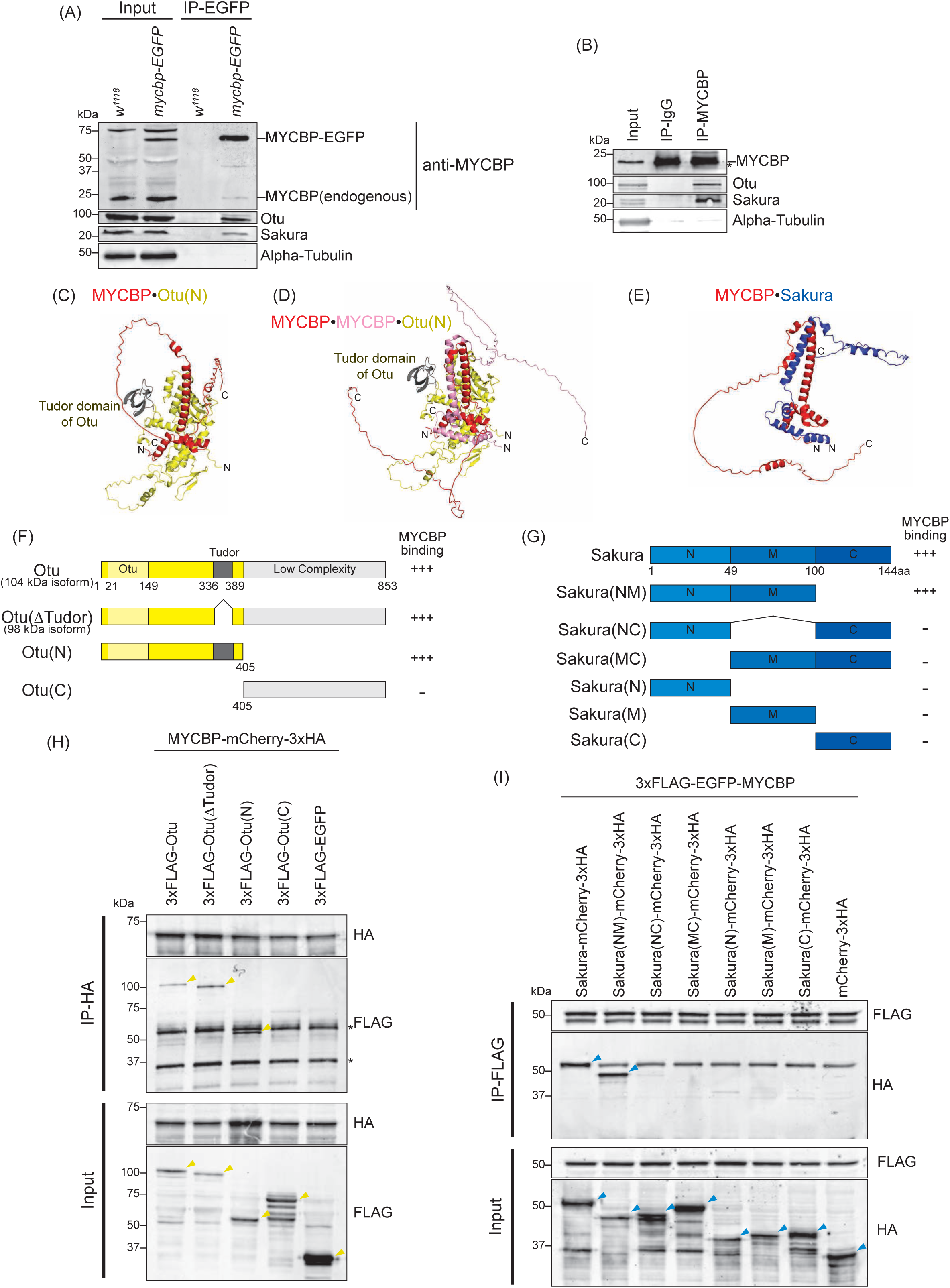
MYCBP interacts with Otu and Sakura. (A) Co-immunoprecipitation (co-IP) using anti-GFP magnetic beads followed by Western blotting. Ovary lysates expressing MYCBP-EGFP in *mycbp^+/+^* background and those from *w^1118^* negative control were analyzed. (B) Co-IP using anti-MYCBP antibodies and ovary lysates from *w^1118^*. IgG was used as a negative control IP. (C-E) Structures of (C) MYCBP•Otu(N), (D) MYCBP•MYCBP•Otu(N), and (E) MYCBP•Sakura predicted using Alphafold. (F, G) Schematic diagrams of full-length and fragment constructs of (F) Otu and (G) Sakura used in co-IP assays. The binding assay results from (H) is summarized. (G) Full-length Sakura and Sakura fragments tested in co-immunoprecipitation assays. N: N-terminal, M: middle, C: C-terminal. The co-IP assay results from (H) and (I) are summarized. (H) Co-IP using anti-HA beads followed by Western blotting. S2 cell lysates co-expressing MYCBP-mCherry-3xHA and 3xFLAG-Otu (full-length or fragments) were tested. 3xFLAG-EGFPserved as a negative control. (I) Co-IP using anti-FLAG beads followed by Western blotting. S2 cell lysates co-expressing 3xFLAG-EGFP-MYCBP and Sakura-mCherry-3xHA (full-length or fragments) were tested. mCherry-3xHA served as a negative control.

Endogenous MYCBP also co-immunoprecipitated with MYCBP-EGFP in ovary lysates, suggesting an interaction between MYCBP proteins (Fig 2A). N-terminally 3xFLAG-EGFP-tagged MYCBP co-immunoprecipitated with C-terminally mCherry-3xHA tagged MYCBP in S2 cells (Fig S1A), confirming the interaction between two MYCBP proteins.

We predicted the structures of the MYCBP•MYCBP, MYCBP•Otu, MYCBP•MYCBP•Otu, and MYCBP•Sakura complexes using AlphaFold (Jumper et al. 2021; Yang et al. 2023), which gave us reasonable predicted structures when using the N-terminal region (1-405aa. Otu(N)), but not the C-terminal region, of Otu (Figs 2C-2E, and S2). In the predicted MYCBP•MYCBP structure, two MYCBP molecule have pseudo-symmetric interaction involving mostly their alpha-helices (Fig S2A). In the predicted structures of both MYCBP•Otu and MYCBP•MYCBP•Otu complexes, Tudor domain of Otu is not involved in the direct interaction with MYCBP. To test these, we created epitope-tagged full-length and truncated Otu— Otu(ΔTudor, corresponding to the endogenous 98 kDa isoform), Otu(N), and Otu(C)—and co-expressed them with epitope-tagged full-length MYCBP in S2 cells. (Fig 2F). Co-immunoprecipitation followed by Western blotting showed that all fragments except Otu(C) interacted with MYCBP (Fig 2H), demonstrating the formation of the MYCBP•Otu and/or MYCBP•MYCBP•Otu complex consistent with the AlphaFold prediction.

AlphaFold predicted a pseudo-symmetric alpha-helical interaction between MYCBP and Sakura, involving the N-terminal and middle regions of Sakura (1-100 aa) (Fig 2E). We generated epitope-tagged full-length and truncated Sakura—Sakura(NM), Sakura(NC), Sakura(MC), Sakura(N), Sakura(M), and Sakura(C) —and tested their ability to bind epitope-tagged full-length MYCBP in S2 cells (Fig 2G). Only full-length Sakura and Sakura(NM) interacted with MYCBP (Fig 2I), supporting the predicted MYCBP•Sakura interaction.

AlphaFold also predicted that MYCBP, Sakura, and Otu could form a ternary complex, MYCBP•Sakura•Otu (Fig 3A). To test this, we performed sequential co-immunoprecipitation from S2 cells co-expressing Sakura-EGFP-HRV3Csite-3xFLAG, MYCBP-mCherry-3xHA, and Myc-Otu (Fig 3B). After FLAG-IP and elution by HRV3C protease cleavage, a second HA-IP detected all three proteins, but not when EGFP-HRV3Csite-3xFLAG was used as a control. Additional co-immunoprecipitation showed that MYCBP-mCherry-3xHA specifically bound Myc-Otu, but not Myc-tagged CycA, Bam, EGFP, or *Drosophila* ortholog of MYC (dMyc) (Fig S1B and S1C), confirming the specificity of the interaction. We conclude that MYCBP, Sakura, and Otu form a ternary complex.

**Fig 3.**
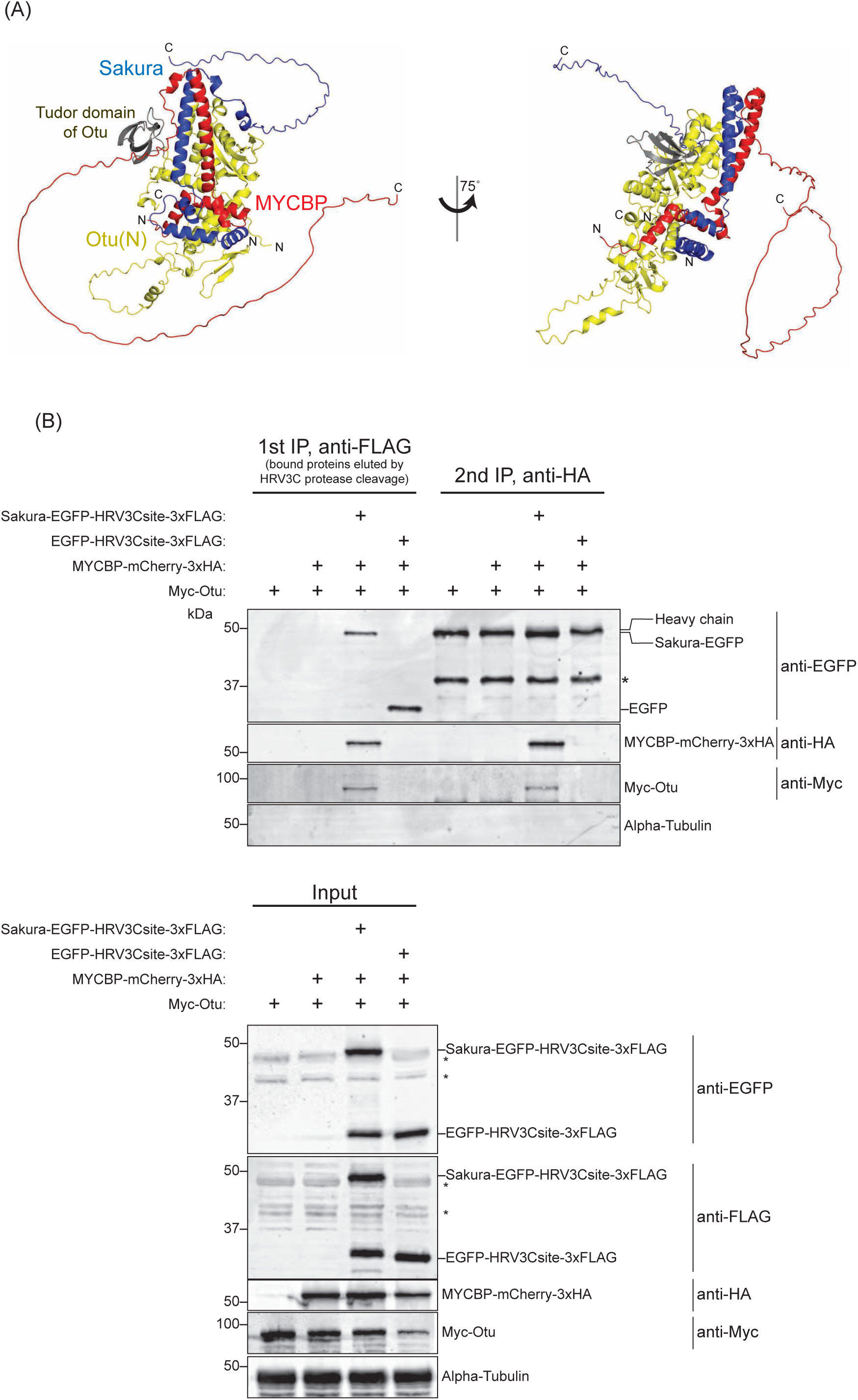
Sequential co-immunoprecipitation shows MYCBP, Sakura, and Otu form a ternary complex. (A) MYCBP•Sakura•Otu structure predicted by AlphaFold. (B) Sequential co-IP followed by Western blotting. S2 cell lysates co-expressing Sakura-EGFP-HRV3Csite-3xFLAG, MYCBP-mCherry-3xHA, and Myc-Otu were used. EGFP-HRV3csite-3xFLAG served as a negative control. The first IP was performed using anti-FLAG beads and eluted using HRV3C protease. Second IP was performed with anti-HA beads. Non-specific bands are marked with *.

Additionally, AlphaFold predicted structures of Sakura•Sakura, Sakura•Otu, and Sakura•Sakura•Otu complexes (Fig S2D-F) and we confirmed the interaction between two Sakura proteins using S2 cells (Fig S1D) and that between Sakura and Otu (Azlan et al. 2024).

### Neither MYCBP nor Sakura affects Otu’s deubiquitinase activity in vitro

Otu possesses deubiquitinase activity (Ji et al. 2017). Our previous study showed that Sakura does not influence Otu’s deubiquitinase activity in vitro (Azlan et al. 2024). Using the same Ub-Rhodamine 110-based assay, we tested whether MYCBP, with or without Sakura, affects Otu’s deubiquitinase activity. The addition of recombinant MYCBP, Sakura, or both had no effect on Otu’s deubiquitinase activity, and neither MYCBP nor Sakura showed deubiquitinase activity (Fig S3).

### *mycbp^null^* mutant flies

To examine MYCBP function *in vivo*, we generated a *mycbp* mutant allele (*mycbp^null^*) by CRISPR/Cas9-mediated deletion of 38 nucleotides (nts) within the MYCBP coding region, resulting in a frameshift and premature stop codon (Fig 1B). The predicted truncated protein consists of 37 N-terminal amino acids of MYCBP followed by a 47-aa frameshifted segment. This fragment is unlikely to be functional and no stable protein product was detected (Fig 4A), indicating a null mutation. Homozygous *mycbp^null/null^* flies were viable, demonstrating that MYCBP is not essential for survival.

**Fig 4.**
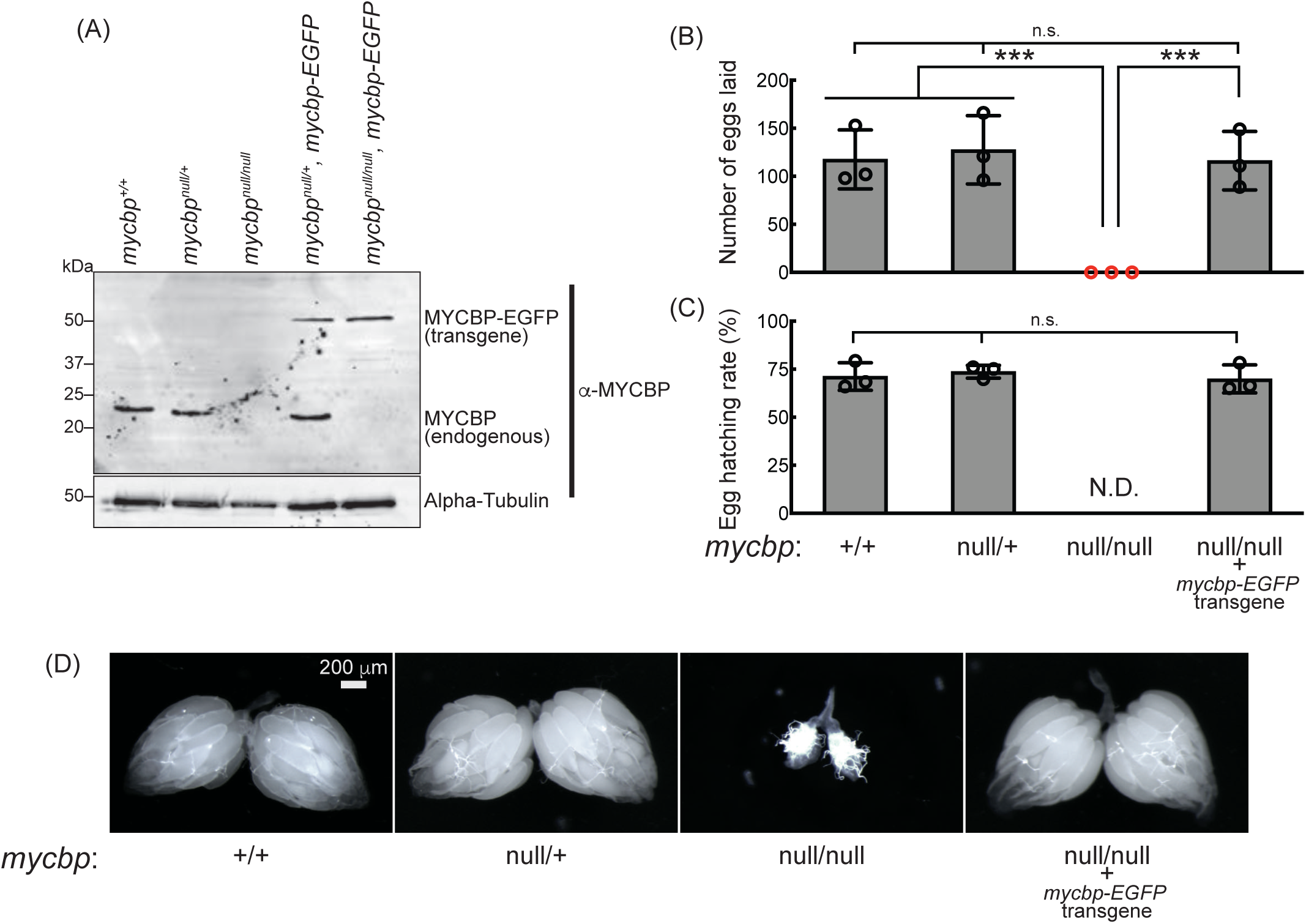
*mycbp^null^* female flies are sterile and have rudimentary ovaries. (A) Western blot of ovary lysates. (B, C) Female fertility assays. (B) The number of eggs laid by test females mated with wild-type (OregonR) males. (C) Hatching rate of the eggs. Mean ± SD (n = 3). P-value < 0.001 (Student’s t-test, unpaired, two-tailed) is indicated by ***. (D) Stereomicroscope images of dissected ovaries. Scale bar: 200 mm.

Western blotting confirmed that MYCBP protein is absent in *mycbp^null/null^* ovaries, but present in wild-type (*mycbp^+/+^*) and heterozygous controls (*mycbp^null/+^*) (Fig 4A). No smaller fragments corresponding to truncated MYCBP were detected in either *sakura^null/+^* or *sakura^null/null^*, suggesting that the fragment is unstable or not expressed.

### MYCBP is essential for female fertility

Given MYCBP’s high expression in ovaries (Fig 1C) and its physical interaction with Otu and (Figs 2 and 3), we hypothesized a role in oogenesis. *mycbp^null/null^* females laid no eggs (Fig 4B and 4C) and their ovaries appeared rudimentary (Fig 4D). Expression of transgenic MYCBP-EGFP in the *mycbp^null/null^* background (*mycbp^null/null^*; *mycbp-EGFP*) fully rescued both fertility and ovary morphology (Fig 4B-4D). Western blotting confirmed that these rescue flies expressed MYCBP-EGFP at levels comparable to endogenous MYCBP in control flies (Fig 4A). MYCBP-EGFP expression in the background of *mycbp*^+/+^ was also detected in testis including germlines such as GSCs (Fig S4A), but *mycbp^null/null^* males were fertile (Fig S4B). Thus, MYCBP is required for female fertility and normal ovary morphology but not for male fertility.

### *mycbp^null/null^* ovaries are germless or tumorous

To investigate the ovary phenotype, we analyzed *mycbp^null/null^* flies expressing a Vasa-EGFP reporter, as Vasa is a known germ cell marker. Some *mycbp^null/null^*ovarioles lacked germ cells ("germless", cyan stars), while others contained germ cells (Fig 5A and 5B). HTS staining revealed overproliferation of GSC-like cells with round spectrosomes in *mycbp^null/null^* ovarioles containing germ cells (Fig 5C, orange stars), indicative of a "tumorous" phenotype as previously described for mutants of *bam*, *otu*, *sxl*, and *sakura* (Smith and King 1966; Gans et al. 1975; Gollin and King 1981; King and Riley 1982; McKearin and Spradling 1990; Bopp et al. 1993; Eliazer et al. 2011; Jin et al. 2013; Yang et al. 2019; Azlan et al. 2024). Additionally, we observed an excess number of cyst cells with branched fusomes that persisted throughout the ovarioles, suggesting abnormal cyst cell development.

**Fig 5.**
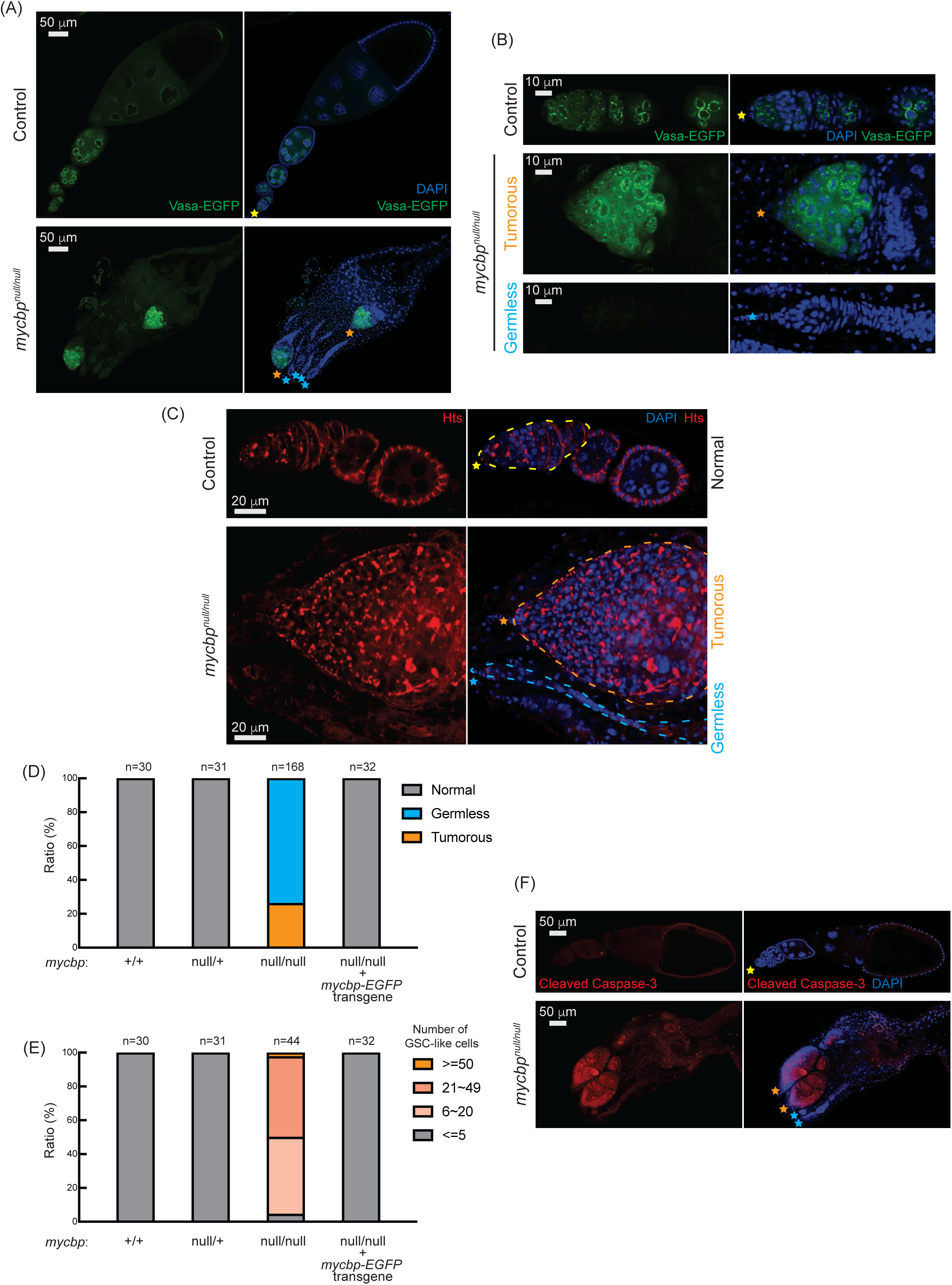
*mycbp^null^* ovaries are germless and tumorous. (A, B) Confocal images of ovaries from control (*mycbp^null/+^*) and *mycbp^null/null^* flies expressing Vasa-EGFP. Vasa-EGFP (green) and DAPI (blue). Yellow, orange, and. cyan stars mark normal ovarioles, tumorous, and germless ovarioles, respectively, in Figure 5. (B) Higher-magnification images of germaria. Scale bars: 50 μm (A), 10 μm (B). (C) Confocal images of ovaries from control (*mycbp^null/+^*) and *mycbp^null/null^* flies stained with anti-Hts to label spectrosomes and fusomes. Hts (red) and DAPI (blue). Germaria are outlined. Scale bars: 20 μm. (D) Percentage of normal, germless, and tumorous ovarioles in indicated genotypes (ages 2-5 days). (E) Quantification of GSC-like cells per germarium (ages 2-5 days). (F) Confocal images of ovaries stained with anti-cleaved Caspase-3. Cleaved caspase-3 (red) and DAPI (blue). Scale bars: 50 μm.

Within the same ovary, both germless and tumorous ovarioles were observed (Fig 5A-5C). 26% of ovarioles were tumorous, while 74% were germless in 2-5-day-old *mycbp^null/null^*flies (n=168) while all ovarioles in control (*mycbp^+/+^* and *mycbp^null/+^*) and *mycbp-EGFP* rescue flies were normal (Fig 5D). The mean number of GSCs or GSC-like cells in 2-5-day-old *mycbp^+/+^*, *mycbp^null/+^*, and *mycbp-EGFP* rescue was 2.0 ± 0.6, 2.2 ± 0.6, and 2.1 ± 0.5, respectively while that for *mycbp^null/null^* was 21.7 ± 12.7 (p-value <0.001) (Fig S5). 95% of *mycbp^null/null^* ovarioles containing germ cells had more than five GSC-like cells (Fig 5E). Tumorous ovarioles showed markedly elevated cleaved Caspase-3 staining, indicating apoptosis (Fig 5F). These results suggest that MYCBP is required for GSC survival, proliferation, and differentiation.

### Loss of *mycbp* results in loss of piRNA-mediated transposon silencing

In control ovaries, Vasa-EGFP localized to the perinuclear nuage in nurse cells (Fig 5A and 5B), which is essential for piwi-interacting RNA (piRNA) biogenesis. This localization persisted in *mycbp^null/null^*tumorous ovarioles, suggesting that MYCBP is not required for Vasa localization.

To assess MYCBP’s role in the piRNA pathway, we used the *Burdock* transposon sensor, which expresses nuclear GFP and β-galactosidase (β-gal) in germ cells but is silenced by piRNAs (Fig 6A) (Handler et al. 2013). In germline-specific *mycbp* RNAi flies (*UAS-Dcr2*, *NGT-Gal4*, *nos-Gal4-VP16* > *mycbp^RNAi^*), we observed strong reporter expression, showing that loss of *mycbp* results in loss of piRNA-mediated transposon silencing.

**Fig 6.**
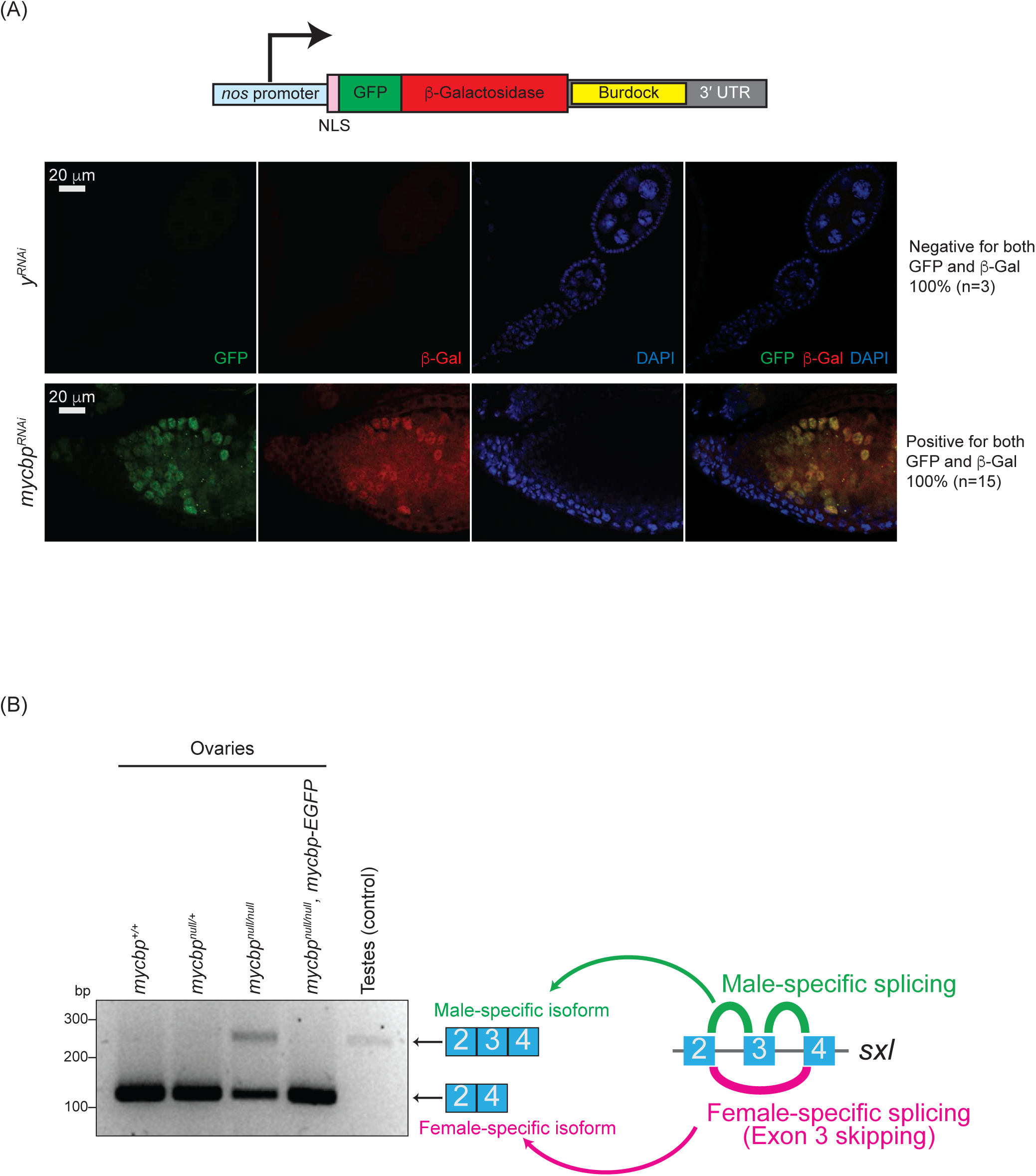
Loss of *mycbp* disrupts piRNA-mediated transposon silencing and *sxl* splicing. (A) Schematic of the *Burdock* sensor construct and representative images of sensor expression in ovaries from control (*y^RNAi^*) and *mycbp^RNAi^*flies. The *Burdock* sensor harbors a *nanos* promoter, a nuclear localization signal (NLS) appended to GFP and β-gal coding sequences, and a target sequence for *Burdock* piRNAs in the 3’UTR. RNAi was driven in the germline using *UAS-Dcr2*, *NGT-Gal4*, and *nos-Gal4-VP16*. GFP (green), β-gal (red), and DAPI (blue). Scale bars: 20 μm. 0/3 control samples showed sensor activation; 15/15 *mycbp^RNAi^* samples did. (B) RT-PCR analysis of *sxl* alternative splicing in ovaries and testes.

### Sex-specific *sxl* mRNA alternative splicing is dysregulated in *mycbp^null/null^* ovaries

Sxl is a master regulator of sex determination in *Drosophila* (Penalva and Sanchez 2003; Salz and Erickson 2010; Grmai et al. 2022). Sex-specific alternative splicing of *sxl* transcripts produces distinct mRNA isoforms: the female-specific, which encodes functional Sxl protein, and the male-specific mRNA isoforms, which does not. In the female germline, loss of Sxl function or disruption of female-specific *sxl* splicing leads to developmental defects, including germline tumors and sterility. Notably, several mutants with tumorous ovariole phenotypes—such as *otu* and *sakura* mutants—exhibit aberrant *sxl* mRNA splicing, resulting in male-specific isoform expression in ovaries and defects in germ cell sexual identity (Bopp et al. 1993; Pauli et al. 1993; Azlan et al. 2024).

As expected, ovaries from control and *mycbp-EGFP* rescue flies expressed only the female-specific *sxl* mRNA isoform, while control testes expressed only the male-specific isoform (Fig 6B). In contrast, *mycbp^null/null^* ovaries showed aberrant expression of the male-specific isoform and a reduced level of the female-specific isoform. These results indicate that female-specific *sxl* splicing is disrupted in the absence of *mycbp*.

### MYCBP is required intrinsically for GSC maintenance and differentiation

To test whether *mycbp* functions autonomously in the germline, we performed mosaic analysis using the FLP-FRT system driven by heat shock-inducible FLP (hs-HLP) (Xu and Rubin 1993; Rubin and Huynh 2015). GSC maintenance was assessed by generating *mycbp^null^* GSC clones and measuring clone loss over time (Xie and Spradling 1998; Yang et al. 2007; Azlan et al. 2024). At four days post-induction, 30.4% of GSCs were marked (GFP-negative) in controls (*FRT82B*) and 20.0% were marked in *mycbp^null^* (*FRT82B*, *mycbp^null^*), establishing initial clone frequencies (Fig 7A). By day 14, marked controle GSCs declined modestly to 19.1% (37.1% loss), whereas marked *mycbp^null^* GSC clones dropped sharply to 4.9% (75.7% loss), indicating that *mycbp* is intrinsically required for GSC maintenance.

**Fig 7.**
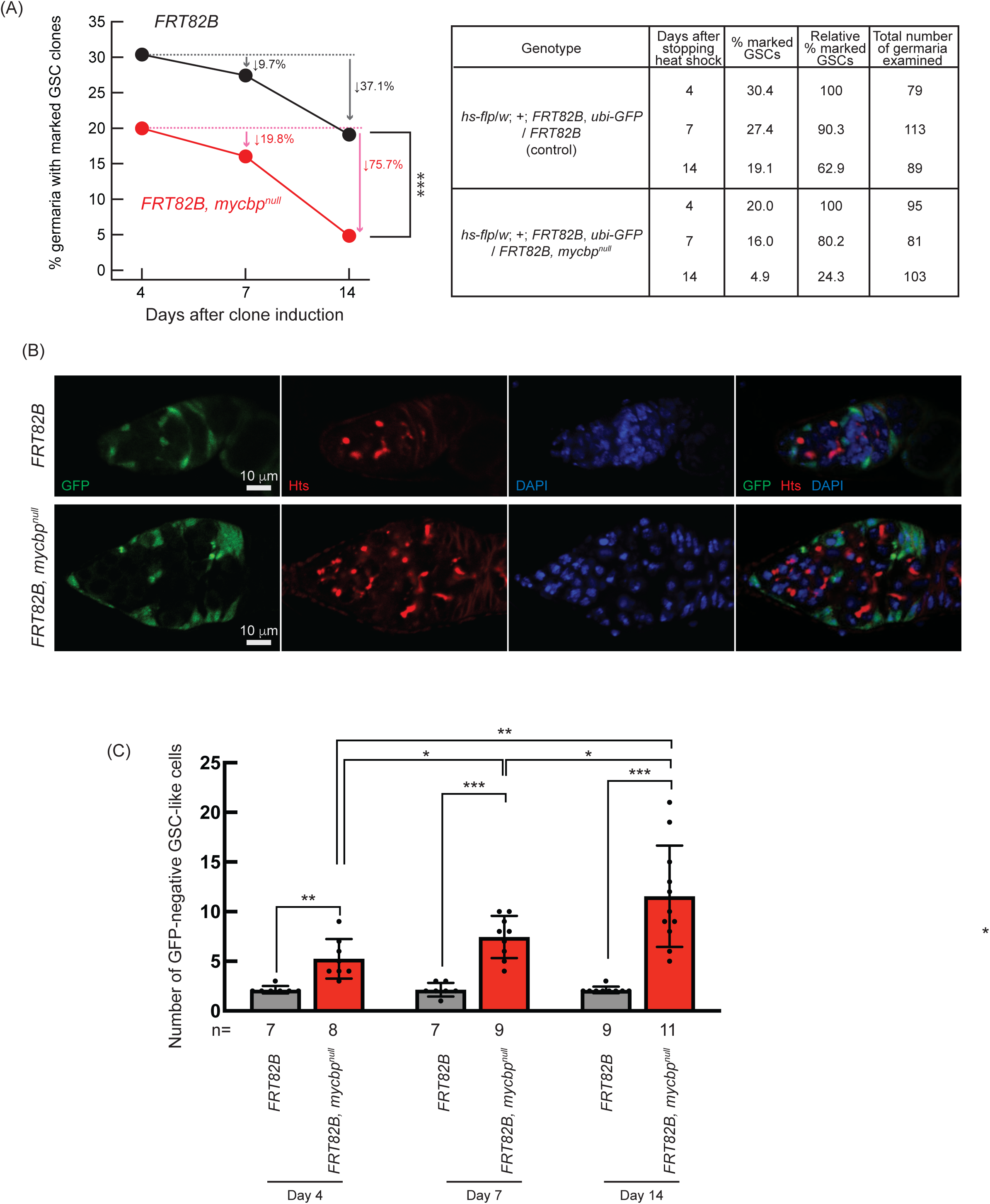
Germline clonal analysis of *mycbp^null^*. (A) Percentage of germaria with marked (GFP-negative) GSC clones at 4, 7, and 14 days after clone induction. P-value < 0.001 (Chi-squared test) is indicated by ***. (B) Confocal images of control and *mycbp^null^* clones. GFP (green), Hts (red), and DAPI (blue). Scale bar: 10 μm. (C) Number of marked (GFP-negative) GSC-like cells per germarium containing marked GSCs at 4, 7, and 14 days after clone induction. GSC-like cells containing round spectrosome were identified by anti-Hts staining. P-value < 0.05, <0.01, and <0.001 indicated by *, **, and *** (Student’s t-test, unpaired, two-tailed).

To determine whether *mycbp* is also intrinsically required for GSC differentiation, we quantified the numbers of marked and unmarked GSC-like cells—germline cells with a round spectrosome—in germaria containing marked GSC clones (Fig 7B). Marked *mycbp^null^*GSC-like cells were significantly more abundant than marked controls at all time points (Fig 7C), and their number increased over time, whereas the number of marked control cells did not (Fig 7C). In contrast, the number of unmarked GSC-like cells in germaria containing marked *mycbp^null^* GSC clones did not differ significantly compared with that in germaria containing marked control GSC clones, and their number did not increase over time (Fig S6). Thus, in the absence of *mycbp*, germline cells exhibit uncontrolled proliferation and tumorous phenotypes. These findings demonstrate that *mycbp* is intrinsically required for both GSCs maintenance and proper differentiation.

### Loss of *mycbp* inhibits Dpp/BMP signaling

The Dpp/BMP signaling pathway plays a central role in regulating GSC self-renewal and differentiation by repressing *bam* in GSCs and de-repressing it in daughter cystoblasts (Fig 8A) (Kirilly and Xie 2007; Hayashi et al. 2020). We hypothesized that the germless and tumorous phenotypes observed in *mycbp^null^* ovaries might result from misregulation of this pathway. To test this, we RNAi-knocked down *mycbp* in the germline using *UAS-Dcr2* and *NGT-Gal4* in the flies carrying the *bam-GFP* reporter (Chen and McKearin 2003b). In controls (*y^RNAi^*), Bam-GFP expression was restricted to 8-cell cysts and absent in later stages (Fig 8B). In contrast, *mycbp* knockdown ovaries exhibited persistent Bam-GFP expression throughout the germarium, including in GSCs (Fig 8B).

**Fig 8.**
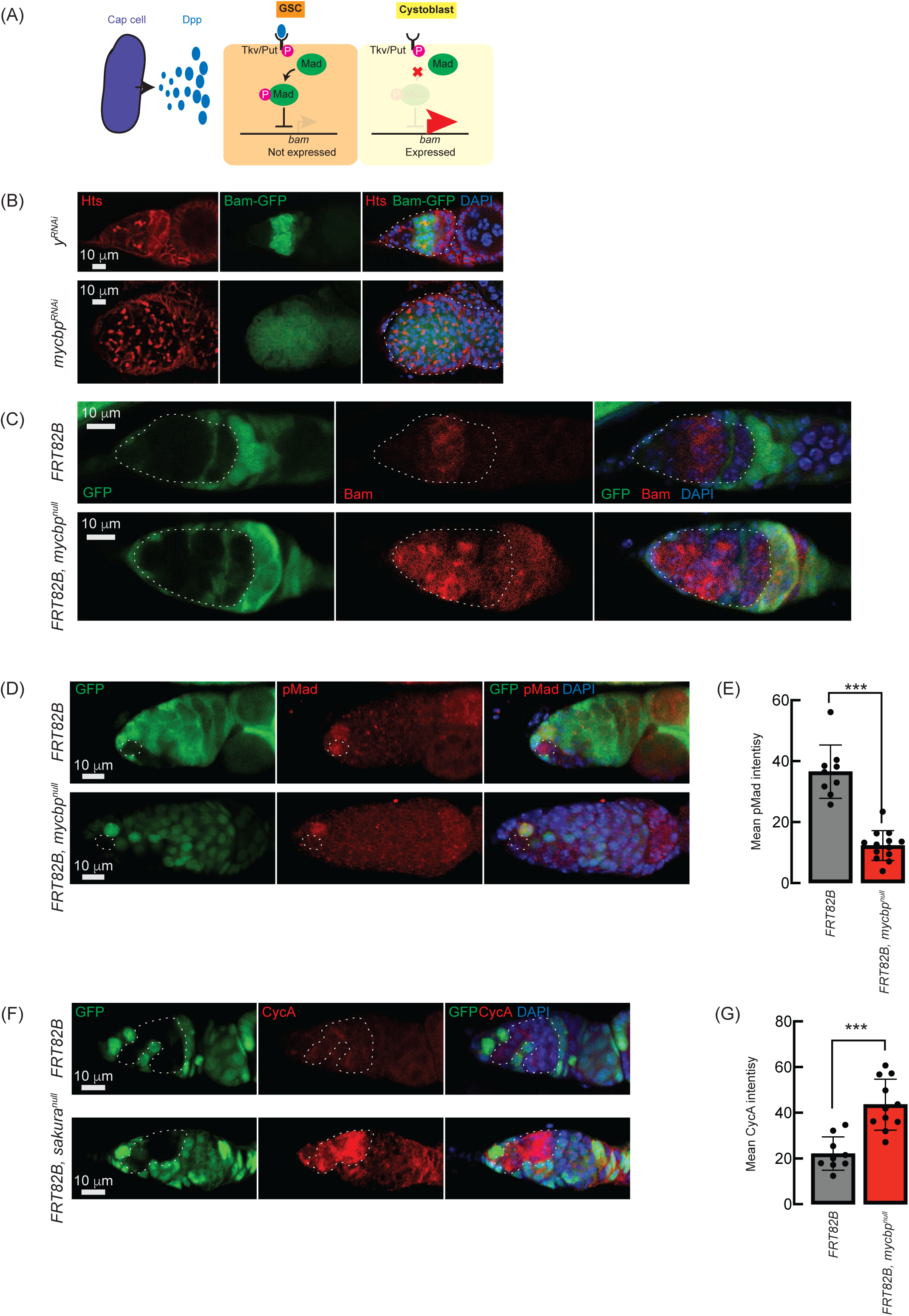
Loss of *sakura* inhibits Dpp/BMP signaling. (A) Schematic of Dpp-mediated *bam* repression via pMad activation. Cap cells secrete diffusible Decapentaplegic (Dpp), which is received by its receptor, a heterodimer of Thick vein (Tkv) and Punt (Put), in GSCs. The activated Dpp signaling eventually phosphorylates Mother-against-dpp (Mad). The phosphorylated Mad (pMad) represses the transcription of *bam*. The repression of *bam* in GSCs is crucial for maintaining their stemness. Cystoblasts do not receive Dpp, and Bam expression is crucial for promoting cystoblast differentiation from GSCs. (B) Confocal images of *bam-GFP* reporter expression in control (*y^RNAi^*) and *mycbp^RNAi^* ovaries (Azlan et al. 2024). RNAi was driven in the female germline using *UAS-Dcr2* and *NGT-Gal4*. Bam-GFP (green), Hts (red), and DAPI (blue). Germaria are outlined. Scale bar: 10 μm. (C, D) Confocal images of germaria with control and *mycbp^null^* germline clones stained with (C) anti-Bam and (D) anti-pMad. GFP (green), Bam/pMad (red), and DAPI (blue). Scale bar: 10 μm. (E) Quantification of pMad intensity in GSC clones. Mean ± SD (n = 9 and 13 for *FRT82B* and *FRT82B, mycbp^null^,* respectively). P-value < 0.001 (Student’s t-test, unpaired, two-tailed) is indicated by ***. (F) Confocal images of germaria with control and *mycbp^null^* germline clones stained with anti-CycA. GFP (green), CycA (red), and DAPI (blue). Scale bar: 10 μm. (G) Quantification of CycA intensity in the germline clones. Mean ± SD (n = 9 and 11 for *FRT82B* and *FRT82B, mycbp^null^*, respectively). P-value < 0.01 (Student’s t-test, unpaired, two-tailed) is indicated by **. Clones in C, D, and F are GFP-negative.

We confirmed this using the FLP-FRT-mediated *mycbp^null^* clones and performed anti-Bam immunostaining. In control clones (*FRT82B*), Bam expression was confined to 8-cell cysts, whereas in *mycbp^null^*clones (*FRT82B*, *mycbp^null^*), Bam was aberrantly expressed throughout the germarium, including in GSCs (Fig 8C). This suggests that in the absence of *mycbp*, Bam expression is no longer repressed by Dpp/BMP signaling in GSCs, resulting in GSC loss, and is no longer shut off after the 16-cell cyst stage.

Dpp/BMP signaling represses *bam* transcription via the transcription factor Mad, which, when phosphorylated (pMad), translocates into the nucleus to repress *bam* transcription (Kirilly and Xie 2007; Kahney et al. 2019; Hinnant et al. 2020). anti-pMad staining revealed significantly reduced pMad in *mycbp^null^* GSC clones compared to control GSCs (Fig 8D and 8E), indicating compromised BMP signaling and transcriptional *bam* de-respression, though Bam is also known to be regulated post-transcriptionally (Pek et al. 2009).

Previous studies have shown that ectopic expression of a stable form of CycA leads to germ cell loss (Chen et al. 2009). This germ cell loss phenotype is also observed upon ectopic *bam* expression in GSCs (Xie and Spradling 1998; Chen and McKearin 2003a; Xia et al. 2010). It was reported that Bam associates with Otu to promote deubiquitination and stabilization of CycA (Ji et al. 2017). Anti-CycA staining showed significantly elevated CycA in *mycbp^null^* clones relative to neighboring wild-type cells in the same germarium (*FRT82B*, *mycbp^null^*) and control clones in control germarium (*FRT82B*) (Fig 8F and 8G), suggesting that Bam misexpression in *mycbp^null^*stabilizes CycA.

### *mycbp* is required for oogenesis in beyond GSCs and germline cysts

Both *mycbp^null/null^* mutation and *mycbp* germline RNAi knockdown using *UAS-Dcr-2* and *NGT-Gal4*, which initiates RNAi in germline cells starting from GSCs, resulted in rudimentary ovaries lacking late-stage germline cells (Fig 4D and 9A), preventing assessment of MYCBP’s role beyond the germarium. However, because MYCBP is expressed throughout egg chamber development, (Fig 1C-E), we hypothesized it may also function in later germline stages.

**Fig 9.**
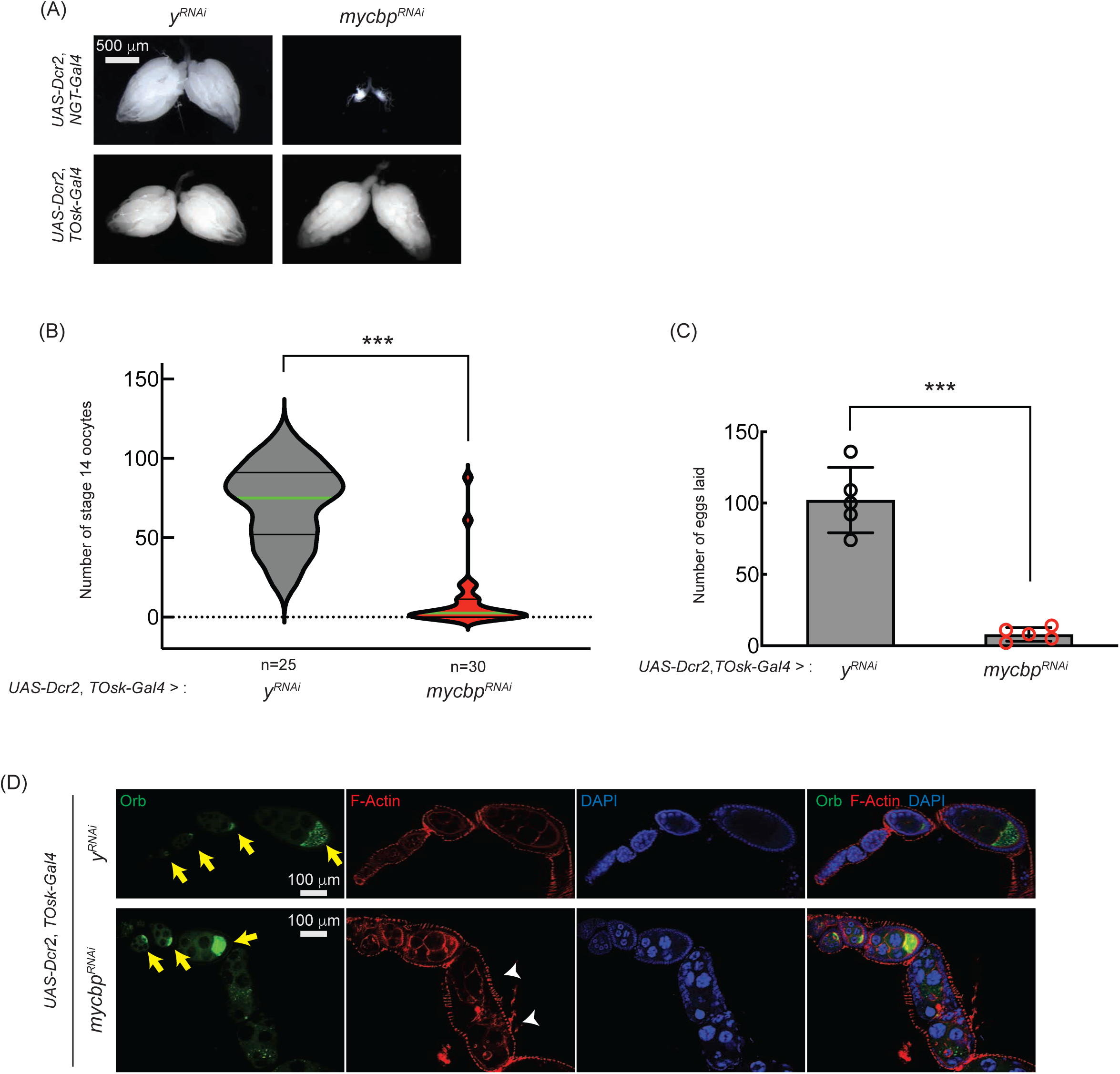
*mycbp* is important for late oogenesis. (A) Stereomicroscope images of dissected ovaries. Scale bar: 500 μm. *y^RNAi^* served as a control. (B, C) Number of stage 14 oocytes per fly (B) and eggs laid (C) in *mycbp* RNAi knockdown usingby *UAS-Dcr2* and *TOsk-Gal4*. Mean ± SD (n = 5). P-value < 0.001 (Student’s t-test, unpaired, two-tailed) is indicated by ***. (D) Confocal images of ovaries stained with phalloidin (F-Actin, red), anti-Orb (Green), and DAPI (blue). Yellow arrows indicate normal Orb localization; white arrowheads indicate cytoskeletal disorganization and loss of Orb localization. Scale bar: 100 μm.

To test this, we used *TOsk-Gal4* (a combination of *osk-Gal4* and *αTub67C-Gal4*) to RNAi-knockdown *mycbp* in germline cells starting from germarium region 2b onward, thereby sparing GSCs and early cysts (ElMaghraby et al. 2022). While *TOsk-Gal4*-driven *mycbp* RNAi did not affect ovary morphology (Fig 9A), it dramatically reduced stage 14 oocyte production and egg laying compared to control RNAi (*y^RNAi^*) (Fig 9B and 9C), suggesting that *mycbp* is important for oogenesis beyond the germline cyst stage.

We also analyzed Oo18 RNA-binding protein (Orb), a marker of oocyte identity. In controls, Orb localized to the posterior within stage ∼4-8 egg chambers (Fig 9D, yellow arrows). In *TOsk-Gal4* > *mycbp^RNAi^* ovaries, Orb localization appeared normal through stage ∼6 (Fig 9D, yellow arrows) but was lost by stage ∼8 with signs of cytoskeletal disorganization (Fig 9D, white arrowheads). These defects likely contribute to the impaired oogenesis observed (Fig 9B and 9C). RNAi efficiency was confirmed by Western blots (Fig 10B and 10C). We conclude that MYCBP plays essential roles in oogenesis beyond early germline stages, similar to Sakura and Otu (Azlan et al. 2024).

**Fig 10.**
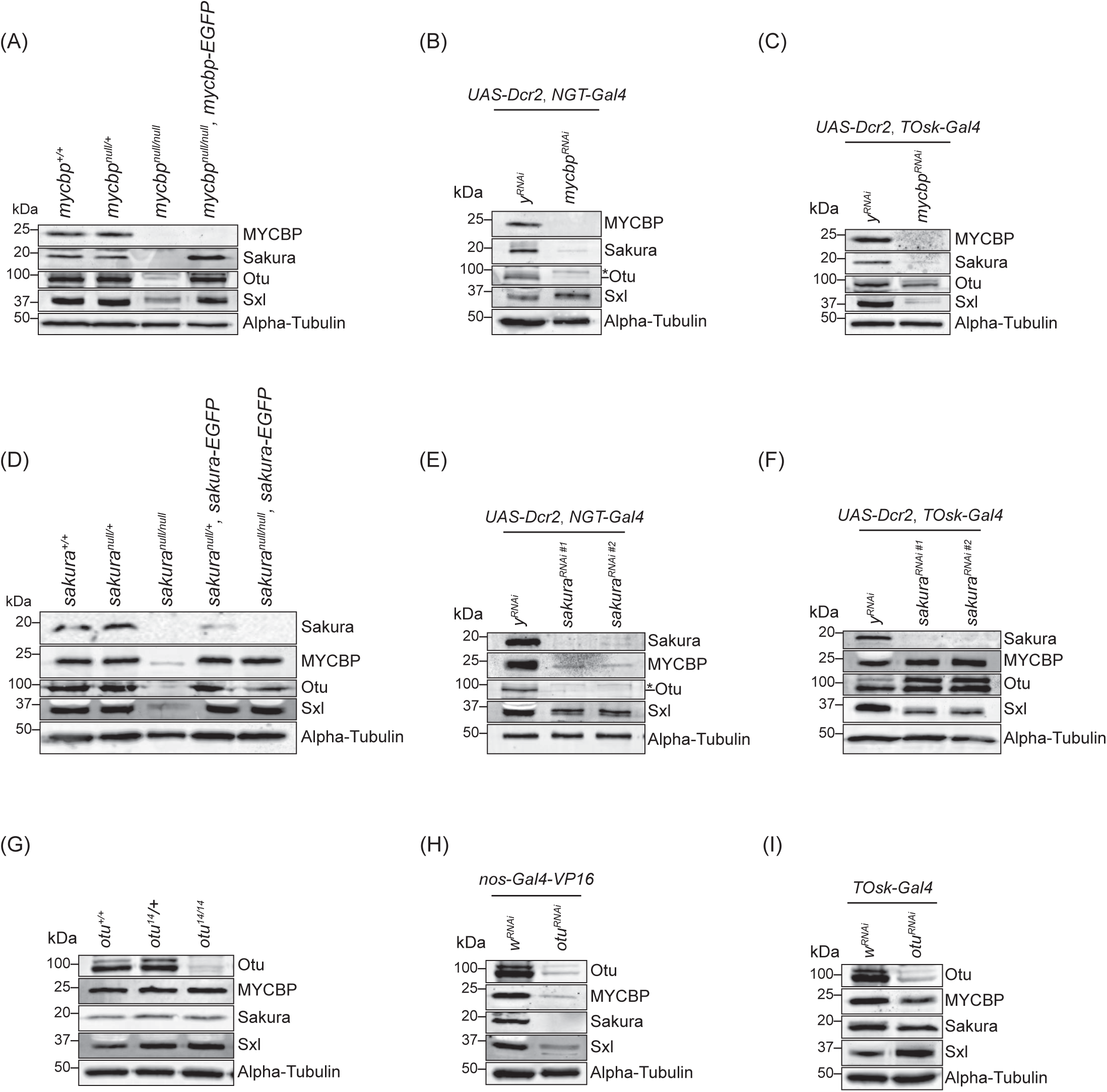
MYCBP is required for Sakura level. Western blot of ovary lysates. The *sakura RNAi* #1 (VDRC: v39727) and #2 (VDRC: v103660). Non-specific bands are marked (*). Sakura and alpha-Tubulin images in (E), Sakura, Otu, and alpha-Tubulin images in (F), Otu and alpha-Tubulin images in (H), Sakura, Otu, and alpha-Tubulin images in (I), are from (Azlan et al. 2024).

### MYCBP is crucial for Sakura levels

Given the formation of the protein complexes among MYCBP, Sakura, and Otu, as well as the shared phenotypes observed in their respective mutants, we investigated whether their protein levels are interdependent. We also assessed Sxl protein levels due to the dysregulation of *sxl* splicing ovserved in their mutants. We performed Western blots using ovary lysates from genetic mutants (*mycbp^null/null^*, *sakura^null/null^*, and *otu^14/14^*) and from RNAi-mediated knockdowns driven by germline-specific Gal4 drivers. Specifically, we used *NGT-Gal4* and *nos-Gal4-VP16* to deplete the proteins starting from GSCs, and *TOsk-Gal4* to deplete proteins from germarium region 2b onward (Fig 10).

In *mycbp^null/null^* ovaries, Sakura and Otu levels were severely reduced, and Sxl levels were moderately reduced (Fig 10A). In *NGT-Gal4 > mycbp^RNAi^* ovaries, Sakura and Otu levels were markedly reduce, while Sxl levels remain unchanged (Fig 10B). In *TOsk-Gal4 > mycbp^RNAi^* ovaries, Sakura and Sxl levels were reduced, but Otu was not affected (Fig 10C). There findings indicate that MYCBP is essential for Sakura levels and may also influence Otu and Sxl levels in a developmental stage-dependent manner.

In *sakura^null/null^* ovaries, MYCBP, Otu, and Sxl levels were severely reduced (Fig 10D). In *NGT-Gal4 > sakura^RNAi^* ovaries, MYCBP and Otu were strongly reduced, but Sxl levels remained unaffected (Fig 10E). In *TOsk-Gal4 > sakura^RNAi^* ovaries, Sxl was reduced, while MYCBP and Otu levels were unchanged (Fig 10F). There data suggest that Sakura may support the expression or stability of MYCBP, Otu, and Sxl proteins in a stage-specific manner.

In *otu^14/14^* ovaries, MYCBP, Sakura and Sxl levels were unchanged (Fig 10G). In contrast, in *nos-Gal4-VP16 > otu^RNAi^* ovaries, all three proteins were reduced (Fig 10H), whereas in *TOsk-Gal4 > otu^RNAi^*ovaries, their levels remained unchanged (Fig 10I). There results suggest that Otu may contribute to the expression and/or stability of MYCBP and Sakura proteins under certain developmental conditions.

It is important to note that whole ovary lysates include both germline and somatic cells. While Sakura and Otu are expressed exclusively in germ cells in ovaries, MYCBP was detectable in both germline and somatic follicle cells (Fig 1E) (Azlan et al. 2024). All Gal4 drivers used (*NGT-Gal4*, *nos-Gal4-VP16* and *TOsk-Gal4*) are germline-specific. Furthermore, ovaries from *mycbp^null/null^*, *sakura^null/null^*, *NGT-Gal4 > mycbp^RNAi^, NGT-Gal4 > sakura^RNAi^*, and *nos-Gal4-VP16 > otu^RNAi^* flies are significantly smaller and degenerated compared to controls (Fig 4D) (Azlan et al. 2024), complicating the interpretation of Western blot results due to tissue loss or altered cell composition. Therefore, we next pursued alternative approaches to more precisely determine their protein level interdependency.

### MYCBP is required for Sakura level and Sakura is important for MYCBP level and its posterior localization within egg chambers

We examined how MYCBP and Sakura influence their protein level and localization within egg chambers each other using FLP-FRT-mediated mosaic analysis. In flies expressing Sakura-EGFP, its signal was markedly reduced in *mycbp^null^*clones (*FRT82B*, *mycbp^null^*) compared to control clones (*FRT82B*), confirming MYCBP is essential for Sakura protein level (Fig 11A).

**Fig 11.**
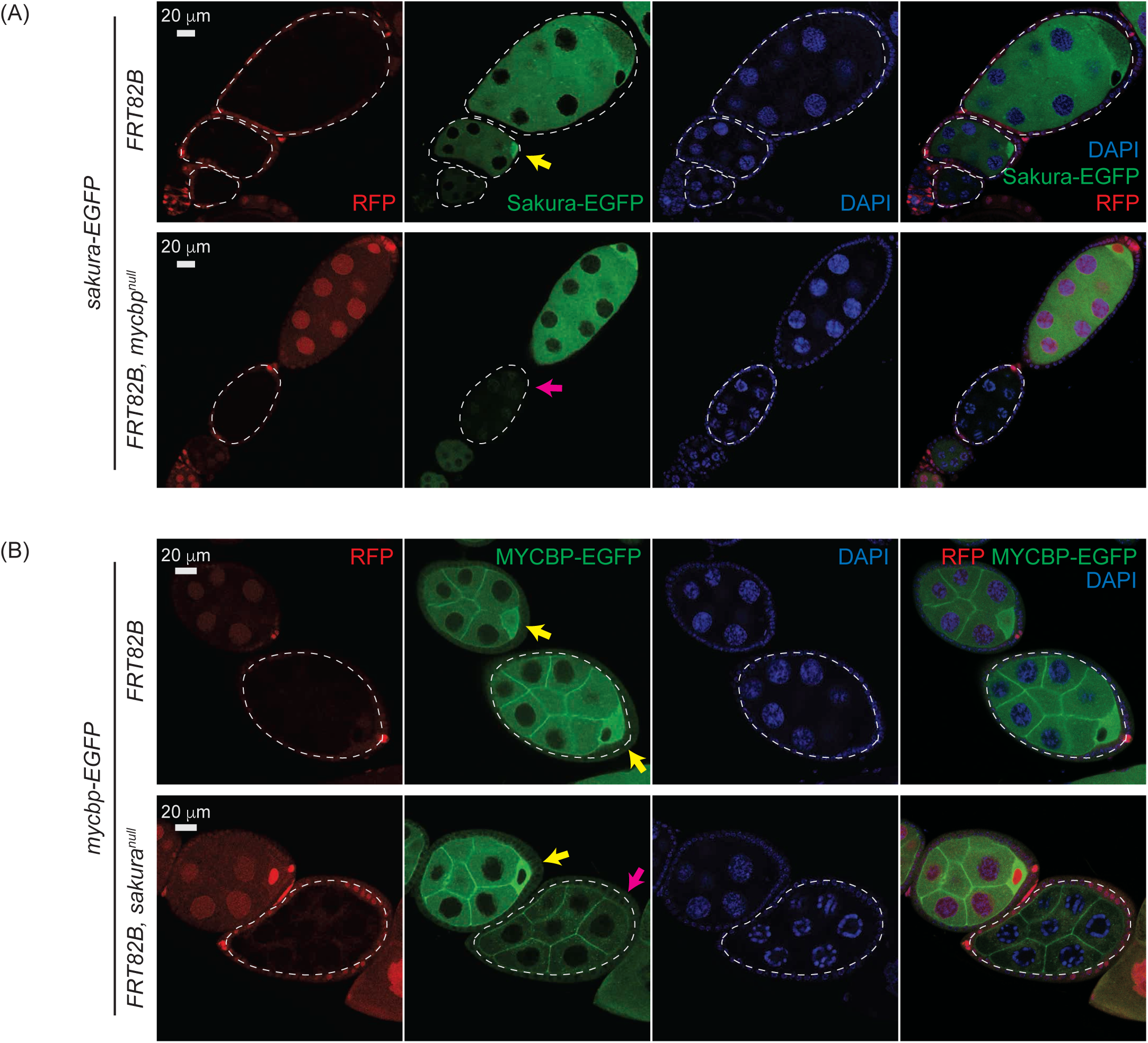
Mutual dependance of MYCBP and Sakura level and localization. (A) Confocal images of egg chambers with control and *mycbp^null^* germline clones expressing Sakura-EGFP. Marked clones (RFP-negative) are outlined. RFP (red), Sakura-EGFP (green), and DAPI (blue). Scale bar: 20 μm. Yellow arrow: normal Sakura-EGFP posterior enrichment; magenta arrow: reduced Sakura-EGFP signal. Fly genotypes used: *hs-flp*/*w*; *sakura-EGFP*/+; *FRT82B*, *ubi-RFP*/*FRT82B*. *hs-flp*/*w*; *sakura-EGFP*/+; *FRT82B*, *ubi-RFP*/*FRT82B*, *mycbp^null^*. (B) Confocal images of egg chambers with control and *sakura^null^* germline clones expressing MYCBP-EGFP. Marked clones (RFP-negative) are outlined. RFP (red), MYCBP-EGFP (green), and DAPI (blue). Scale bar: 20 μm. Yellow arrow: normal MYCBP-EGFP posterior enrichment; magenta arrow: reduced levels and loss of posterior localization of MYCBP-EGFP. Fly genotypes used: *hs-flp*/*w*; *mycbp-EGFP*/+; *FRT82B*, *ubi-RFP*/*FRT82B*. *hs-flp*/*w*; *mycbp-EGFP*/+; *FRT82B*, *ubi-RFP*/*FRT82B*, *sakura^null^*.

Conversely, in flies expressing MYCBP-EGFP, its level was decreased and its posterior localization was lost in *sakura^null^* clones (*FRT82B*, *sakura^null^*) compared in control clones (*FRT82B*) (Fig 11B), indicating that Sakura is required for MYCBP level and its posterior localization within egg chambers.

### MYCBP and Sakura are required for Otu localization to developing oocytes

Finally, we examined Otu level and localization within egg chambers using *otu-EGFP* and *otu(ΔTudor)-EGFP* transgenes in control, *mycbp^null^*, and *sakura^null^* clones. We also stained for Orb, which is localized to developing oocytes. Otu-EGFP and Otu(ΔTudor)-EGFP levels were similar among all genotypes (Fig 12), suggesting MYCBP and Sakura do not affect Otu levels. However, Otu-EGFP and Otu(ΔTudor)-EGFP failed to localize to the posterior within the egg chambers in *mycbp^null^* and *sakura^null^* clones. Importantly, Orb remained properly localized in these clones, suggesting that oocyte specification was intact. Thus, MYCBP and Sakura dispensable for Otu level but are required for its proper localization to developing oocytes.

**Fig 12.**
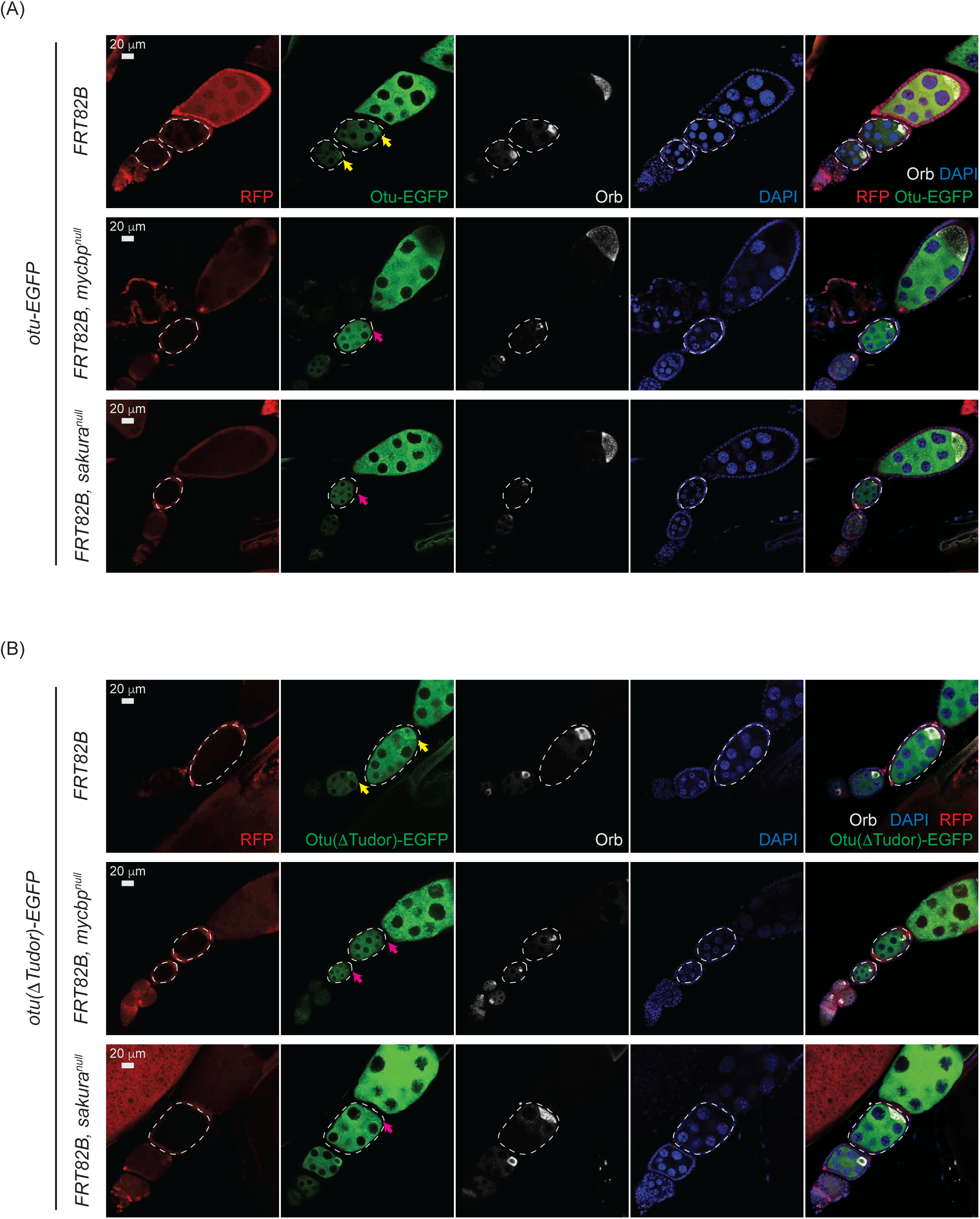
Otu localization to the developing oocyte depends on MYCBP and Sakura. Confocal images of egg chambers with control, *mycbp^null^*, and *sakura^null^* germline clones expressing (A) Otu-EGFP or (B) Otu(∆Tudor)-EGFP. Marked clones (RFP-negative) are outlined. RFP (red), Otu-EGFP or Otu(∆Tudor)-EGFP (green), Orb (white), and DAPI (blue). Scale bar: 20 μm. Yellow arrows: normal posterior enrichment of Otu-EGFP and Otu(∆Tudor)-EGFP; magenta arrows: loss of localization. Orb localization remains intact in mutant clones. Fly genotypes used: *hs-flp*/*w*; *otu-EGFP*/+; *FRT82B*, *ubi-RFP*/*FRT82B*. *hs-flp*/*w*; *otu-EGFP*/+; *FRT82B*, *ubi-RFP*/*FRT82B*, *mycbp^null^*. *hs-flp*/*w*; *otu-EGFP*/+; *FRT82B*, *ubi-RFP*/*FRT82B*, *sakura^null^*. *hs-flp*/*w*; *otu(∆Tudor)-EGFP*/+; *FRT82B*, *ubi-RFP*/*FRT82B*. *hs-flp*/*w*; *otu(∆Tudor)-EGFP* /+; *FRT82B*, *ubi-RFP*/*FRT82B*, *mycbp^null^*. *hs-flp*/*w*; *otu(∆Tudor)-EGFP* /+; *FRT82B*, *ubi-RFP*/*FRT82B*, *sakura^null^*.

## Discussion

We previously showed that Sakura (also known as Bourbon (Mercer et al. 2025)) and Otu form a protein complex (Azlan et al. 2024). In this study, we identified MYCBP, encoded by the previously uncharacterized gene *CG17202*, as a binding partner of both Otu and Sakura. Our data support that MYCBP binds with itself, Sakura, and Otu, forming binary and ternary complexes, including the MYCBP•Sakura•Otu ternary complex. Structural predictions suggest that MYCBP and Sakura resemble each other and engage in a pseudo-symmetric interaction. Mutations in *mycbp*, *otu*, and *sakura* result in strikingly similar phenotypes, and all three proteins are highly expressed in germline cells of the ovary, localize to the cytoplasm, and are enriched in developing oocytes. These observations strongly indicate that MYCBP, Sakura, and Otu function cooperatively in the germline during oogenesis.

Since there are multiple protein association states that can be possibly formed among MYCBP, Sakura, and Otu, including MYCBP alone, Sakura alone, Otu alone, MYCBP•MYCBP, Sakura•Sakura, MYCBP•Sakura, MYCBP•Otu, MYCBP•MYCBP•Otu, Sakura•Otu, Sakura•Sakura•Otu, and MYCBP•Sakura•Otu, the relative expression levels among the three proteins and potential regulatory mechanisms for their interaction may determine the relative abundance of these multiple protein complexes, which could be critically important for GSC maintenance and differentiation and oogenesis.

In *mycbp^null^* germline clone cells, Sakura protein levels were severely reduced, and Otu lost its localization to developing oocytes, despite unchanged Otu protein levels and normal posterior localization of Orb (Fig 11A and 12). These results indicate that MYCBP is required for Sakura protein expression and/or stability, as well as for proper Otu localization. Similarly, in *sakura^null^*germline clones, MYCBP levels were reduced, and both MYCBP and Otu lost their posterior localization, again without affecting Otu levels or Orb localization (Fig 11B and 12), indicating that Sakura is crucial for MYCBP protein expression and/or stability, as well as for proper MYCBP and Otu localization to developing oocytes. These mutual dependencies of protein expression/stability and oocyte localization among MYCBP, Sakura, and Otu further support the model that they function as protein complexes.

Although MYCBP and Sakura did not directly affect Otu’s deubiquitinase activity in vitro using Ub-Rhodamine 110 as a model substrate (Fig S3), this does not rule out the possibility that they influence Otu’s enzymatic activity in vivo. For instance, they may modulate Otu’s substrate specificity. Previous work has shown that Otu also interacts with Bam—primarily through its Otu domain—to form a deubiquitinase complex that deubiquitinates and thereby stabilizes CycA, promoting GSC differentiation (Ji et al. 2017). It is possible that MYCBP and Sakura regulate the interaction between Otu and Bam and/or modulate the enzymatic activity of the Otu•Bam complex. For example, binding of MYCBP and/or Sukura to Otu may be mutually exclusive with Bam binding. Alternatively, MYCBP, Sakura, Otu, and Bam might form a ternary complex. Further studies are required to elucidate whether and how MYCBP and Sakura influence Otu’s protein interactions and enzymatic function.

Otu also functions as an RNA-binding protein, and its deubiquitinase activity is enhanced by RNA binding (Ji et al. 2019). Sxl controls both alternative mRNA splicing and translation of downstream targets, and promotes its own expression via a positive autoregulatory loop (Inoue et al. 1990; Bell et al. 1991; Valcarcel et al. 1993; Chau et al. 2012). We and others have shown that female-specific splicing of *sxl* mRNA is disrupted in *mycbp*, *sakura* and *otu* mutant ovaries, leading to production of the male-specific isoform (Fig 6B) (Bopp et al. 1993; Azlan et al. 2024). Bam, together with Bgcn, Mei-P26, and Sxl, binds *nanos* mRNA—a key stem cell maintenance— and represses its translation after germ cells exit the niche (Wang and Lin 2004; Li et al. 2009; Chau et al. 2012; Li et al. 2013). Identifying the RNA targets and deubiquitinase substrates of Otu beyond CycA and how MYCBP and Sakura regulate these Otu’s activities will be critical to understanding their roles in oogenesis and other developmental processes. MYCBP•Sakura•Otu complex may bind directly to RNAs and regulate post-transcriptional processes such as *sxl* alternative splicing and translational control of oogenic RNAs.

Sakura is exclusively expressed in female germline cells including GSCs (Azlan et al. 2024), and MYCBP is also highly expressed in these cells (Fig 1). However, MYCBP is additionally expressed at lower levels in somatic follicle cells in egg chambers and in other tissues, including testes, and Otu is broadly expressed in various tissues such as the gut and testis as well as in female germline cells in ovaries (Steinhauer and Kalfayan 1992). These differential expression patterns suggest that Otu may have tissue-specific functions depending on the presence or absence of MYCBP and Sakura.

Transposons pose significant threat to genomic stability by inducing DNA damage if not properly silenced (Levin and Moran 2011). piRNAs suppress transposons through transcriptional and post-transcriptional silencing mechanisms, thus preserving genome integrity (Huang et al. 2017; Yamashiro and Siomi 2018). Loss of piRNA function results in transposon derepression, increased DNA damage, germ cell apoptosis, arrested oogenesis, and sterility (Kang et al. 2018; Moon et al. 2018). Because damaged germ cells can transmit harmful mutations to the next generation, selective elimination of defective germ cells is critical for maintaining germline integrity of a species (Chu et al. 2014; Ota and Kobayashi 2020). We found that loss of function of *mycbp*, *sakura or otu* impairs piRNA-mediated transposon silencing (Fig 6A) and causes apoptosis (Fig 5F) (Azlan et al. 2024). Thus, the germless phenotypes may arise, at least in part, from activation of a transposon-induced apoptotic elimination program. It will be important to investigate whether MYCBP, Sakura, and Otu’s have any direct roles in the piRNA pathway.

MYCBP and Otu are conserved through human (human MYCBP, also known as AMY-1, and OTUD4) while Sakura is not. Human MYCBP was suggested to bind via its C-termina region to the N-terminal region of C-MYC and stimulate the activation of E-box-dependent transcription by C-MYC. (Taira et al. 1998). However, we showed that *Drosophila* MYCBP does not bind *Drosophila* ortholog of MYC (dMyc) (Fig S1C). Alphafold suggested that human MYCBP and OTUD4 form complexes, MYCBP•OTUD4 and/or MYCBP•MYCBP•OTUD4, via the N-termina region of OTUD4 (Fig S2I and S2J), suggesting the evolutionary conserved interaction between MYCBP ortholog and Otu ortholog.

This study identifies and characterizes evolutionary conserved MYCBP as a novel, essential regulator of oogenesis in *Drosophila*. Together with Sakura and Otu, MYCBP likely controls germline cell fate decision, maintenance, and differentiation.

## Materials and Methods

### Fly strains

We generated the *mycbp^null^*strain by introducing indels into the MYCBP coding region using the CRISPR/Cas9 genome editing, as previously reported (Zhu et al. 2018a; Zhu et al. 2018b; Zhu et al. 2019b; Zhu and Fukunaga 2021; Azlan et al. 2024; Taira et al. 2025). The transgenic *mycbp*-*EGFP* strain was established following previously published methods (Fukunaga et al. 2012; Kandasamy and Fukunaga 2016; Zhu and Fukunaga 2021). A DNA fragment containing the MYCBP coding sequence flanked by ∼1.2 kb upstream and ∼1 kb downstream genomic sequences was cloned. EGFP was fused in-frame to the C-terminus of the MYCBP coding sequence. The construct was inserted into a pattB plasmid and integrated at the 25C6 landing site using the attP40 fly line and the PhiC31 integrase system (BestGene).

Transgenic *otu-EGFP, otu(ΔTudor)-EGFP,* and *sakura-EGFP* strains were previously described (Azlan et al. 2024). The *mycbp-RNAi* (VDRC: v41628), *y-RNAi* (v106068), *TOsk-Gal4* (v 314033), Burdock sensor [*UAS-Dcr2; NGT-Gal4; nosGal4-VP16, nos>NLS_GF’_lacZ_vas-3’UTR_burdock-target*] (v 313217) strains were from Vienna *Drosophila* Resource Center. *UAS-Dcr-2; NGT-Gal4* (BDSC: 25751) and *FRT82B/TM6C, Sb* (BDSC: 86313) were obtained from the Bloomington Stock Center. The *bam-GFP* reporter (DGRC: 118177) and *vasa-EGFP* knocked-in fly (DGRC: 118616) were from Kyoto *Drosophila* Stock Center. *hsFLP*; *FRT82B*, *ubi-RFP*/*TM6B* strain was a kind gift from Dr. Wu-Min Deng (Tulane University).

### Fertility assay

Fertility assays were performed as described previously (Zhu et al. 2018b; Liao et al. 2019; Zhu et al. 2019a; Zhu and Fukunaga 2021; Azlan et al. 2024; Taira et al. 2025). For female fertility, five virgin females of the test genotype were crossed with three wild-type (OregonR) males in cages containing 6-cm grape juice agar plates supplemented with wet yeast paste at 25°C. Plates were replaced daily. Eggs laid on the third plate (from day 3 to day 4) were counted, and hatching was assessed after an additional 24 horus of incubation at 25°C. At least three cages per genotype were analyzed.

For male fertility, a single test male was mated with five wild-type (OregonR) virgin females in a vial at 25°C. After 3 days, the females were transferred to a new vial (vial 1), and then to new vials every 2 days for a total of four vials. Females were removed after 2 days in the fourth vial. Total progeny from all vials were counted. At least five males per genotype were tested.

### MYCBP antibody generation

Recombinant full-length MYCBP with a C-terminal HRV3Csite-6xHis tag was expressed in *E. coli* using a modified pET vector (Fukunaga and Doudna 2009) and purified via Ni-sepharose (Cytiva). The His-tag was removed via HRV3C protease cleavage, and the protein was further purified on a HiTrapQ HP column (Cytiva). This antigen was used to generate rabbit polyclonal anti-sera (Pocono Rabbit Farm & Laboratory, Inc.). Rabbit polyclonal anti-MYCBP antibodies were affinity-purified using recombinant MYCBP-HRV3Csite-6xHis protein conjugated to Affigel-15 (Bio-rad).

### Immunostaining

Stereomicroscope images of dissected ovaries were taken using a Leica M125 stereomicrocsope. Ovaries from 2- to 5-day-old, yeast-fed females were dissected in 1x PBS (137 mM NaCl, 2.7 mM KCl, 10 mM Na_2_HPO_4_, 1.8 mM KH_2_PO_4_, pH 7.4) and fixed for 30 minutes at room temperature in a fix buffer (4% formaldehyde, 15 mM PIPES (pH 7.0), 80 mM KCl, 20 mM NaCl_2_, 2 mM EDTA, and 0.5 mM EGTA). Samples were washed in PBX (1x PBS + 0.1% Triton X-100) and blocked in a blocking buffer (PBX with 2% donkey serum, 3% BSA [w/v], and 0.02% NAN_3_ [w/v]) for 1 hour at room temperature. The ovaries were then incubated with primary antibodies diluted in the blocking buffer overnight at 4°C. Samples were washed three times with PBX and incubated with Alexa Fluor-conjugated secondary antibodies for 2 hours at room temperature, washed again, and mounted in VECTASHIELD® PLUS antifade mounting medium with DAPI (H-2000, Vector lab). Confocal images were acquired on a Zeiss LSM700 confocal microscope at the Johns Hopkins University School of Medicine Microscope Facility.

The primary antibodies used for immunostaining were mouse anti-HTS (1B1) (DSHB, AB_528070, dilution: 1/100), mouse anti-Bam (DSHB, AB_10570327, 1/20), mouse anti-CycA (DSHB, AB_528188, 1/100), mouse anti-Orb (DSHB, AB_528419, 1/100), rabbit anti-pMad (Cell Signaling, Phospho-SMAD1/5 (Ser463/465) mAb #9516, 1/200), and rabbit anti-cleaved caspase-3 (Cell Signaling, Cleaved Caspase-3 (Asp175) #9661, 1/200). Secondary antibodies used were Alexa Fluor 488 Donkey anti-Mouse Igg (ThermoFisher, A21202, 1/100), Alexa Fluor 594 Donkey anti-Mouse Igg (ThermoFisher, A21203, 1/100), Alexa Fluor 594 Donkey anti-Rabbit Igg (ThermoFisher, A21207, 1/100), and Alexa Fluor 647 Donkey anti-mouse Igg (ThermoFisher, A31571, 1/100). Rhodamine phalloidin (ThermoFisher, R415, 1/100) was used to stain F-Actin.

### Germline clonal analysis

*mycbp^null^* germline clones were generated using FLP/FRT-mediated recombination (Rubin and Huynh 2015). *mycbp^null^* GSC clones were induced by heat-shocking 3-day-old females of the genotype *hs-flp*/*w*; *+; FRT82B, ubi-GFP/FRT82B, mycbp^null^* at 37°C for 1 hour, twice daily with an 8-hour interval. Controls were *hs-flp*/*w*; +; *FRT82B, ubi-GFP/FRT82B.* Ovaries were dissected 4, 7, and 14 days post-heat shock.

To generate *mycbp^null^* mutant clones in the presence of transgenic reports, files of following genotypes were used. *hs-flp*/*w*; *otu-EGFP*/+; *FRT82B*, *ubi-RFP*/*FRT82B*, *mycbp^null^*, *hs-flp*/*w*; *otu(ýTudor)-EGFP* /+; *FRT82B*, *ubi-RFP*/*FRT82B*, *mycbp^null^* and *hs-flp*/*w*; *sakura-EGFP*/+; *FRT82B*, *ubi-RFP*/*FRT82B*, *mycbp^null^*. Control genotypes lacking the *mycbp^null^* allele were also analyzed. Flies were dissected 3-5 days after clone induction.

### Western blot

Lysates of hand-dissected ovaries and tissues were prepared by homogenizing in RIPA buffer (50 mM Tris-HCl [pH 7.4], 150 mM NaCl, 1% [v/v] IGEPAL CA-630, 0.1% [w/v] sodium dodecyl sulfate (SDS), 0.5% [w/v] sodium deoxycholate, 1 mM ethylenediaminetetraacetic acid (EDTA), 5 mM dithiothreitol, and 0.5 mM phenylmethylsulfonyl fluoride (PMSF)) (Kandasamy et al. 2017; Zhu et al. 2018b; Azlan et al. 2024; Taira et al. 2025). Homogenates were centrifuged at 21,000g at 4°C for 10 min, and the protein concentrations of the supernatant were determined using the BCA protein assay kit (Pierce). Fifteen μg of total protein was loaded per lane for Western blot.

The sources and dilutions of the primary antibodies were as below. Rabbit anti-MYCBP (1/10000, generated in this study), Rabbit anti-Sakura (1/10000, (Azlan et al. 2024)), rabbit anti-Otu (1/10000, (Azlan et al. 2024)), mouse anti-Sxl M18 (1/1000, DSHB, AB_528464), mouse anti-Sxl M114(1/1000, DSHB, AB_528463), rabbit anti-alpha-Tubulin [EP1332Y] (1/10000, Abcam, ab52866), mouse anti-alpha-Tubulin [12G10] (1/10000, DSHB, AB_1157911), mouse anti-FLAG (1/10000, Sigma, F1804), mouse anti-HA (1/10000, Sigma, H3663), and mouse anti-GFP [GF28R] (1/3000, Invitrogen, 14-6674-82). IRDye 800CW goat anti-mouse IgG (LiCor), IRDye 800CW goat anti-rabbit IgG (LiCor), IRDye 680RD goat anti-mouse (LiCor), and IgG IRDye 680RD goat anti-rabbit (LiCor) were used as secondary antibodies. The membranes were scanned using the Li-Cor Odyssey CLx Imaging System.

### Mass spectrometry

Immunoprecipitation of MYCBP-EGFP protein was performed using the GFP-Trap Magnetic Agarose Kit (Proteintech, gtmak-20) on dissected ovaries from flies harboring *mycbp-EGFP* transgene, with *w1118* flies as controls. Ovaries were homogenized in 200 μL ice-cold lysis buffer (10 mM Tris-HCl [pH 7.5], 150 mM NaCl, 0.5 mM EDTA, 0.05% [v/v] IGEPAL CA-630) containing 1× protease inhibitor cocktail (100× protease inhibitor cocktail contains 120 mg/ml 1 mM 4-(2-aminoethyl) benzene sulfonyl fluoride hydrochloride (AEBSF), 1 mg/ml aprotinin, 7 mg/ml bestatin, 1.8 mg/ml E-64, and 2.4 mg/ml leupeptin). After homogenization, the tubes were placed on ice for 30 minutes, and the homogenates were extensively pipetted every 10 minutes. The lysates were then centrifuged at 17,000x g for 10 minutes at 4°C. The supernatants were transferred to pre-chilled tubes, and 300 μL dilution buffer (10 mM Tris/Cl pH 7.5, 150 mM NaCl, 0.5 mM EDTA) supplemented with 1x protease inhibitor cocktail were added. The diluted lysates were then added to the GFP-trap magnetic beads in 1.5 mL tubes and rotated for 1 hour at 4°C. After separating the beads with a magnetic tube rack, the beads were washed three times with 500 μL wash buffer (10 mM Tris/Cl pH 7.5, 150 mM NaCl, 0.05 % [v/v] IGEPAL CA-630). Proteins were eluted with 40 μL acidic elution buffer (200 mM glycine pH 2.5) followed by immediate neutralization with 5 uL neutralization buffer (1 M Tris pH 10.4).

As a quality control before mass spectrometry, ∼5 μL of the samples were mixed with an equal volume of 2× SDS-PAGE loading buffer (80 mM Tris-HCl [pH 6.8], 2% [w/v] SDS, 10% [v/v] glycerol, 0.0006% [w/v] bromophenol blue, 2% [v/v] 2-mercaptoethanol), heated at 95 °C for 3 min, and run on 4–20% Mini-PROTEAN® TGX™ Precast Protein Gels (Biorad, #4561094). Silver staining was then performed by using Pierce™ Silver Stain Kit (ThermoFisher, 24612) to assess the quality of the immunoprecipitated protein samples. Mass spectrometry was conducted at the Mass Spectrometry Core at the Department of Biological Chemistry, Johns Hopkins School of Medicine, as previously described (Zhu and Fukunaga 2021; Azlan et al. 2024).

### Co-immunoprecipitation

For endogenous MYCBP co-IP, ovaries from wilt-type (*w^1118^*) flies were homogenized in ice-cold lysis buffer (10 mM Tris-HCl [pH 7.5], 150 mM NaCl, 0.5 mM EDTA, 0.05% [v/v] IGEPAL CA-630) containing 1× protease inhibitor cocktail. Lysates were centrifuged at 21,000g at 4°C for 10 min, and supernatants were used for immunoprecipitation. Four μg of rabbit anti-MYCBP and control rabbit IgG (Cell Signaling, #2729) were incubated with 50 μL of Dynabeads Protein G (ThermoFisher, 10004D) for 20 min at room temperature. The beads were washed once with PBST (1x PBS with 0.1% Tween-20), then incubated with lysates at room temperature for 30 min, followed by three washes with PBST. Proteins were eluted with 2x SDS-PAGE loading buffer (80 mM Tris-HCl [pH 6.8], 2% [w/v] SDS, 10% [v/v] glycerol, 0.0006% [w/v] bromophenol blue, 2% [v/v] 2-mercaptoethanol) and heated at 70°C for 10 min. After bead separation using a magnetic tube rack, the eluted proteins in 2x SDS-PAGE loading buffer were heated again at 95°C for 3 min.

For transient expression in S2 cells, a total of 1 μg plasmid DNA (pAc5.1/V5-HisB. Invitrogen) was transfected into cells in 6-well plates using Effectene (Qiagen, 301425). After 3 days, cells were harvested and lysed in ice-cold lysis buffer with 1× protease inhibitor cocktail, and centrifuged at 17,000x g for 10 min at 4°C. For HA-IP, supernatants were incubated with 25 μL (0.25 mg) of Pierce™ Anti-HA Magnetic Beads (ThermoFisher, 88837) at room temperature for 30 min. The beads were then washed three times with TBST (1x TBS [50 mM Tris/HCl and 150 mM NaCl, pH 7.6] with 0.05% Tween-20). For FLAG-IP, 2 μg of mouse anti-FLAG (Sigma, F1804) was pre-bound to 50 μL of Dynabeads Protein G (ThermoFisher, 10004D) for 10 min at room temperature. The beads were washed once with PBST. The S2 cell lysate supernatant was incubated with the washed beads at room temperature for 15 min. The beads were washed three times with PBST. In both anti-HA and anti-FLAG immunoprecipitations, proteins were eluted with 2× SDS-PAGE loading buffer. For anti-HA, the beads in 2x SDS-PAGE loading buffer were heated at 95°C for 7 min. For anti-FLAG, the beads were heated at 70°C for 10 min, then were separated using a magnetic tube rack. The eluted proteins in 2x SDS-PAGE loading buffer were heated again at 95°C for 3 min.

### Sequential co-immunoprecipitation

S2 cells were transfected with the Sakura-EGFP-HRV3Csite-3xFLAG, EGFP-HRV3Csite-3xFLAG, MYCB-mCherry-3xHA, and Myc-Otu plasmid constructs (pAc5.1/V5-HisB. Invitrogen) with the Effectene. A total of 1 μg plasmids were transfected in each well of 6-well plates. After 3 days, cells were harvested, lysed and centrifuged as described above.

For the first IP, 2 μg of mouse anti-FLAG (Sigma, F1804) was pre-bound to 50 μL of Dynabeads Protein G and was incubated with lysate supernatant at room temperature for 15 min. Beads were washed three times with PBST, then proteins were eluted by incubation with 100 μL cleavage buffer (25 mM Tris-HCl pH 7.4, 150 mM NaCl, 5% glycerol, 2 mM EDTA) containing 25 nM GST-HRV3C protease at 4°C for 6 hours. 20 μL of the eluate was mixed with 2x SDS-PAGE loading buffer and heated at 95°C for Western blot analysis.

The remaining ∼80 μL eluate was subjected to a second IP using 25 μL (0.25 mg) of Pierce™ Anti-HA Magnetic Beads (ThermoFisher, 88837) pre-equilibrated in lysis buffer. After a 30 min incubation at room temperature, beads were washed three times with TBST and proteins were eluted by heating at 95°C for 7 min in 2× SDS-PAGE loading buffer.

### RT-PCR

Total RNAs from ovaries and testes and were prepared using miRVana (Thermo Fisher Scientific), followed by DNase treatment with Turbo DNase (Thermo Fisher Scientific). cDNA was synthesized from 1 μg of RNA using SuperScript^TM^ VILO^TM^ MasterMix (Thermo Fisher Scientific). To examine *sxl* alternative splicing, PCR was performed using GoTaq Green Master Mix (Promega) with primers *sxl*-F (5′-CTCACCTTCGATCGAGGGTGTA-3′) and *sxl*-R (5′-GATGGCAGAGAATGGGAC-3′) followed by agarose gel electrophoresis and SYBR Safe staining.

### In vitro deubiquitination assay

Purified Otu, luciferase, Sakura (Azlan et al. 2024), and MCYBP proteins (latter purified as for antibody generation) were mixed with Ub-Rhodamine 110 (Ubiquitin-Proteasome Biotechnologies, M3020) in a 30 μL in reaction (20 mM Tris-HCl, pH 7.5, 200 mM NaCl, 5 mM MgCl_2_, 2 mM DTT). Reactions were performed for 60 min in a black 384-well low-volume plate and fluorescence signals were measured using SpectraMax i3x Multi-Mode microplate reader (excitation/emission: 485/20 nm and 530/20 nm).

## Acknowledgments

We thank Dr. Wu-Ming Deng, Bloomington Drosophila Stock Center, Vienna Drosophila Resource Center, and Kyoto Drosophila Stock Center for fly strain stocks. We thank the Johns Hopkins University School of Medicine Microscope Facility for use of the Zeiss LSM700, supported by NIH grant S10OD016374 awarded to Dr. Scot C. Kuo. We thank Ms. Lauren DeVine, Dr. Josh Smith, Dr. Bob Cole, and Dr. Chan-Hyun Na at the Johns Hopkins University School of Medicine Mass Spectrometry and Proteomics Core Facility for the mass-spec analysis. This work was supported by the grants from the National Institutes of Health [R35GM145352 and R03AI178064] and Johns Hopkins University Catalyst Award to RF.

## Author Contributions

Conceptualization, Methodology, Investigation and Writing, A.A. and R.F.; Funding Acquisition and Supervision, R.F.

## Supplemental Figure Legends

**Fig S1.**
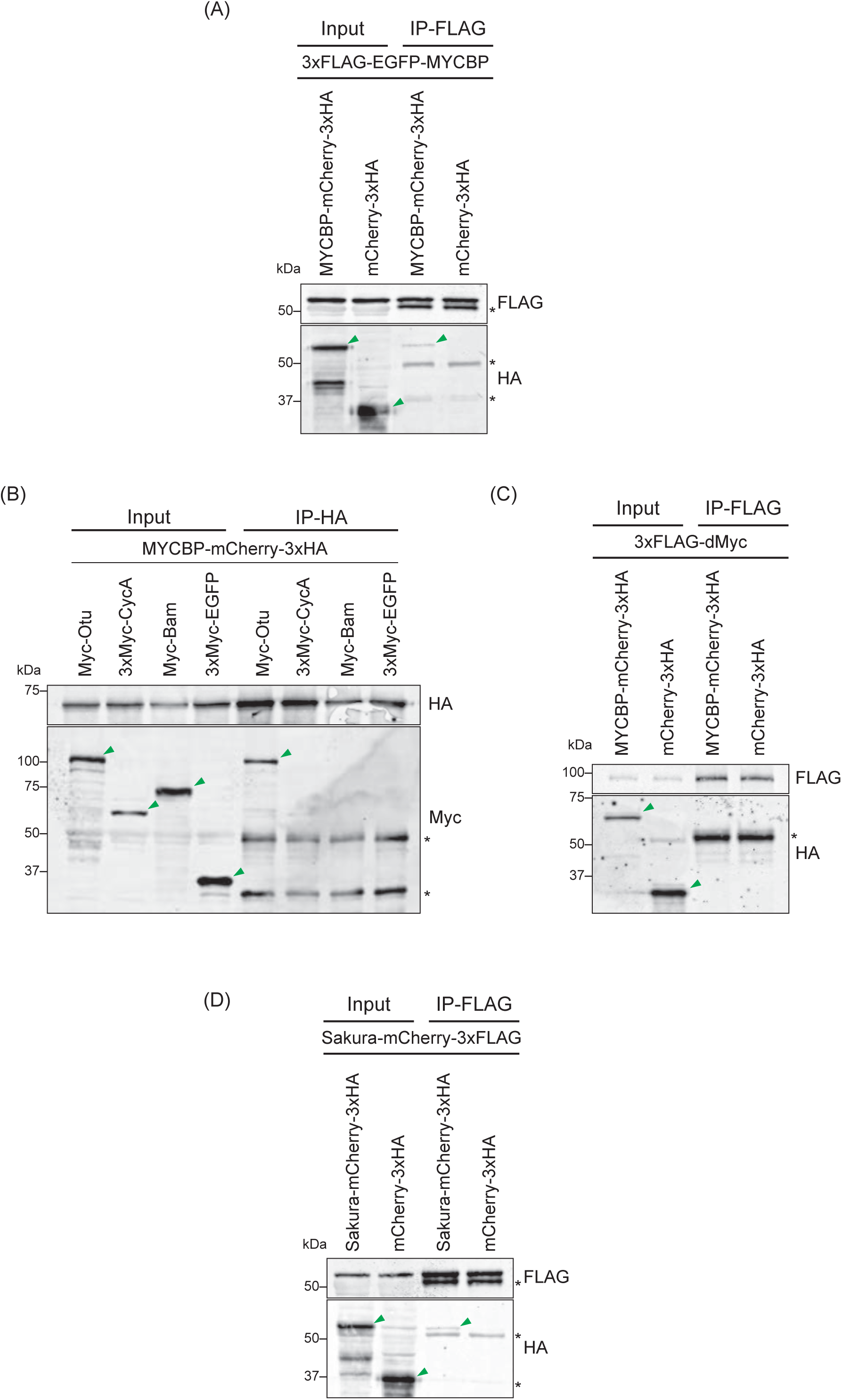
MYCBP and Sakura form homodimers and MYCBP binds Myc-Otu, but not 3xMyc-CycA, Myc-Bam, 3xMyc-EGFP, or dMyc. Co-immunoprecipitation using S2 cell lysates and anti-FLAG (A, C, D) or anti-HA beads (D) followed by Western blotting.

**Fig S2.**
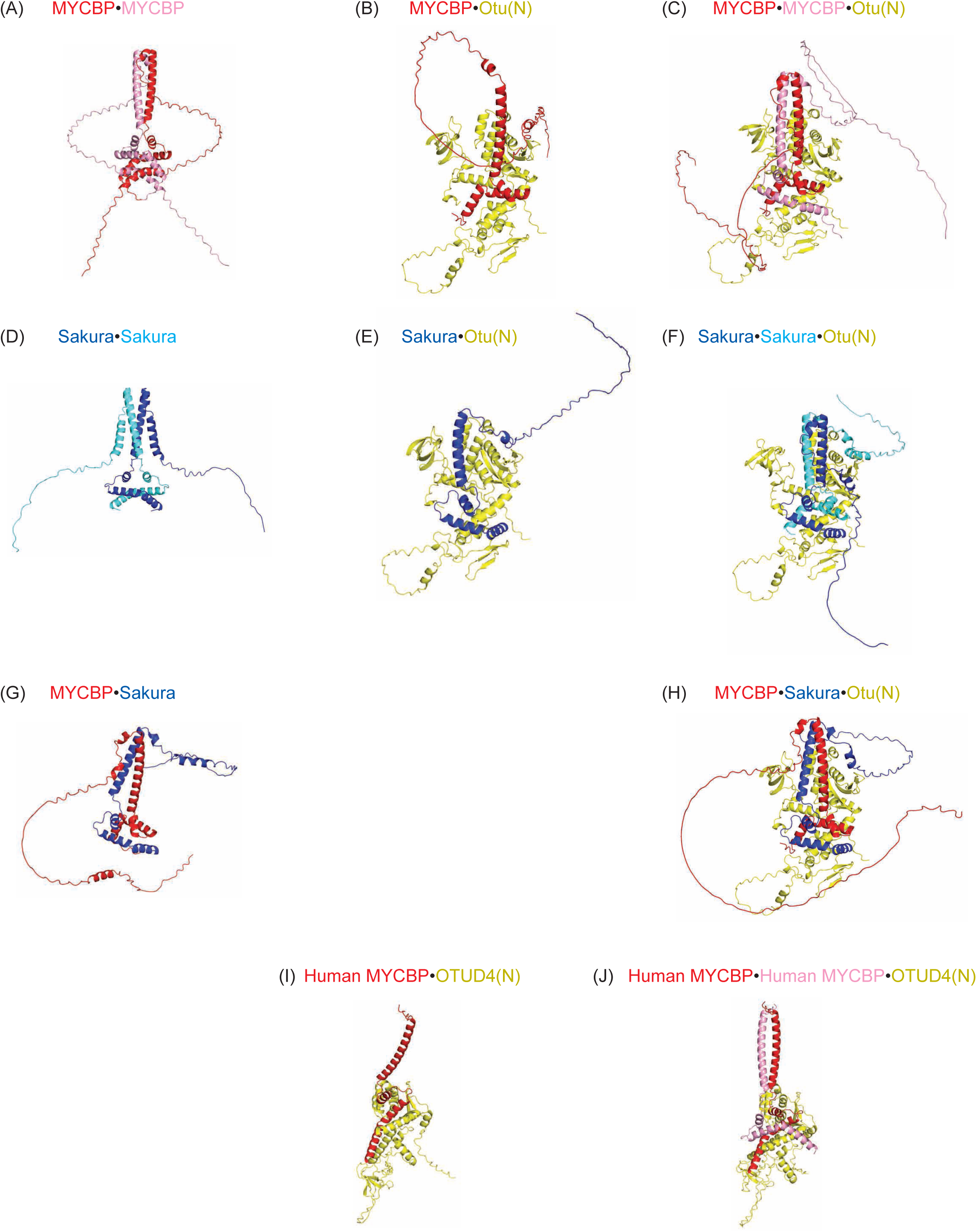
Alphafold-predicted structures. Protein complex structures predicted using Alpha-fold. full-length MYCBP, Sakura, and human MYCBP are used while Otu(N) is 1-405 aa region and OTUD4(N) is 1–350 aa region.

**Fig S3.**
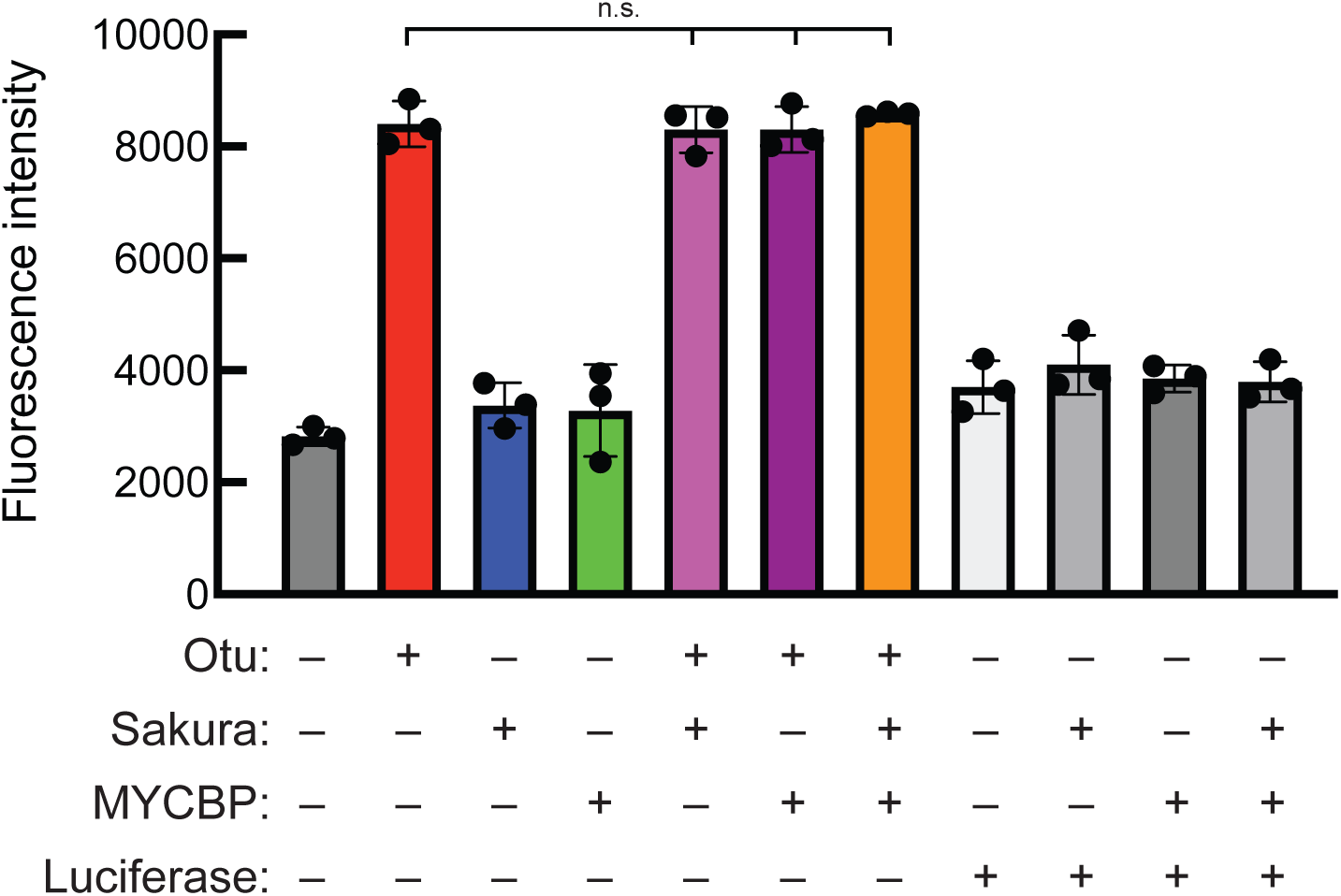
In vitro deubiquitination assay. Fluorescence intensity. Mean ± SD (n = 3). Firefly Luciferase was used as a negative control.

**Fig S4.**
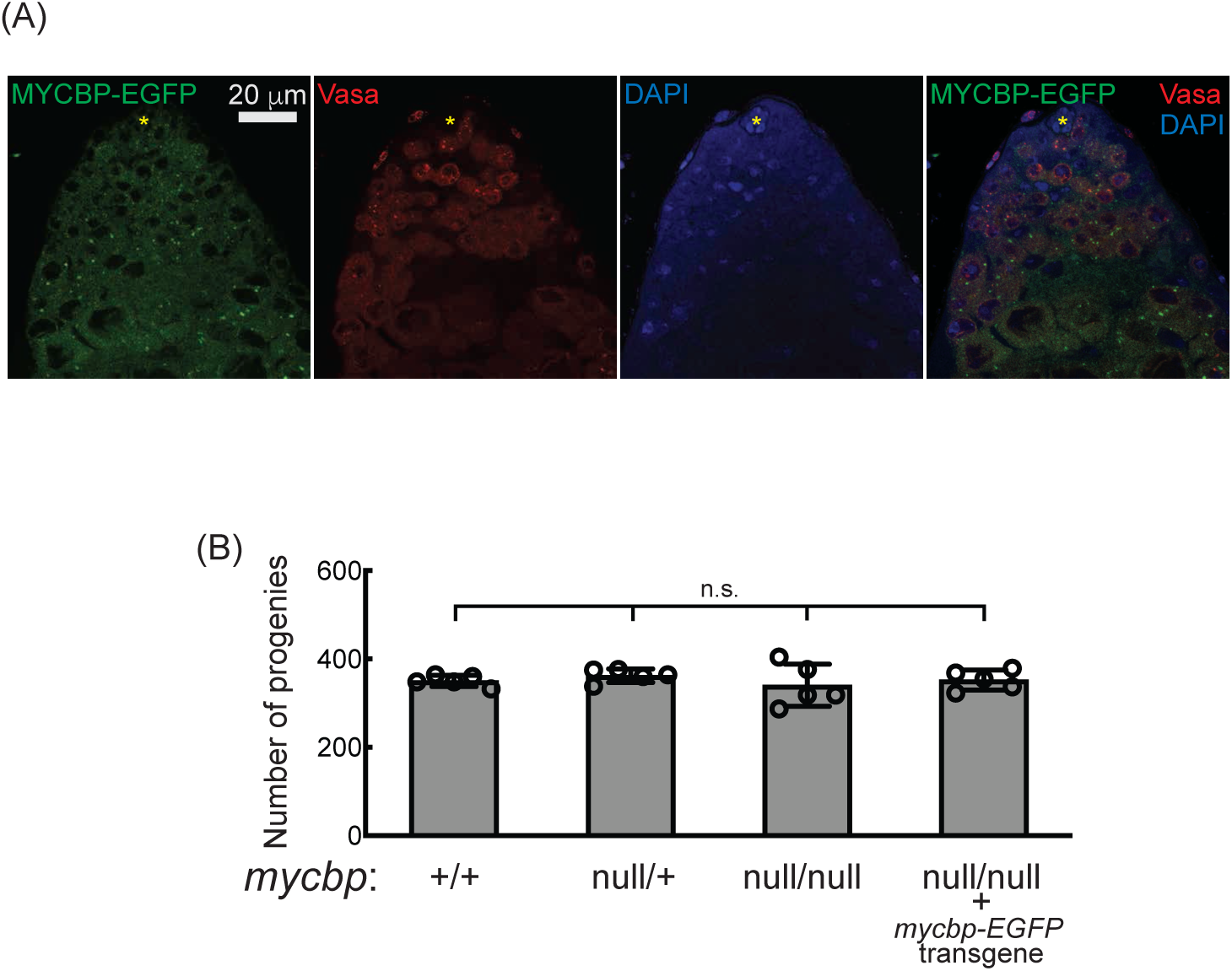
Male fertility assay. (A) Confocal images of the apical tip region of a testis from a *mycbp-EGFP* transgenic fly. MYCBP-EGFP (green), Vasa (red), and DAPI (blue). Hub cells are marked with yellow star. Scale bar: 20 μm. (B) Numbers of progeny obtained from crosses between test males and wild-type (OregonR) virgin females. Mean ± SD (n = 5).

**Fig S5.**
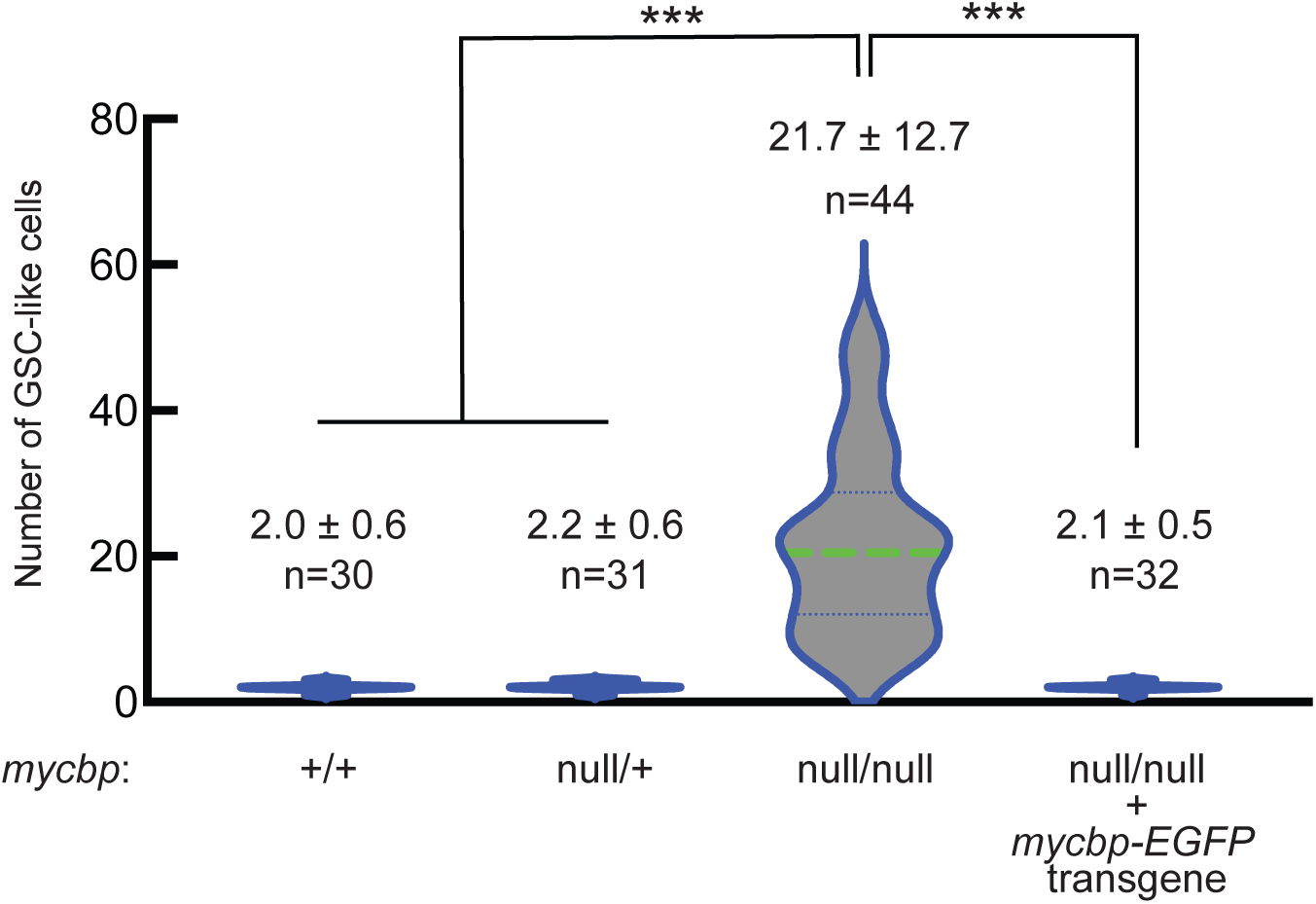
*mycbp^null^* ovaries are tumorous. Violin plots showing the number of GSC-like cells per germaria in 2-5-day-old flies. Mean ± SD. P-value < 0.001 (Student’s t-test, unpaired, two-tailed) is indicated by ***.

**Fig S6.**
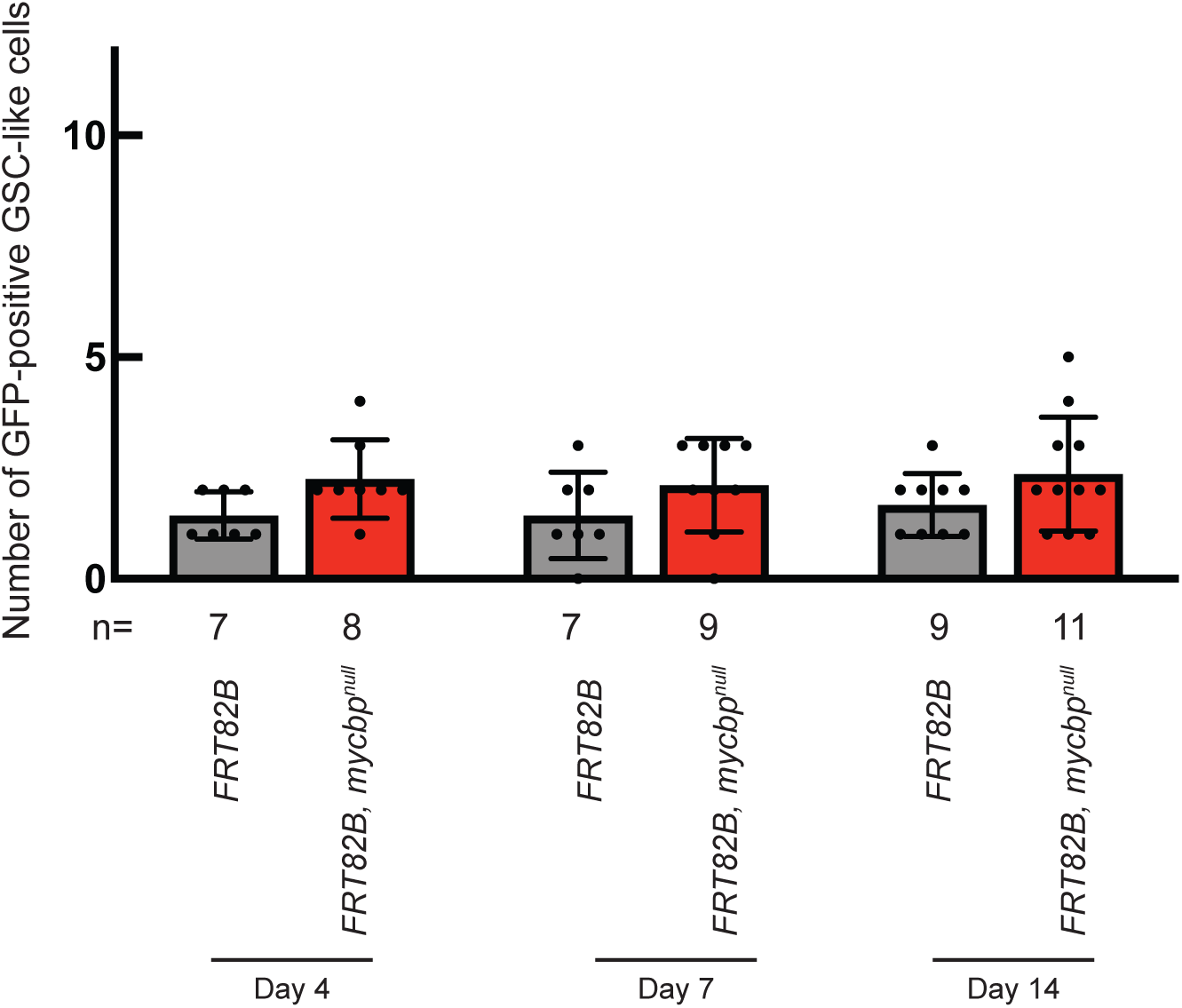
*mycbp^null^* germline clones intrinsically cause tumorous phenotypes. Number of unmarked (GFP-positive) GSC-like cells in germaria containing marked (GFP-negative) GSCs at 4, 7, and 14 days after clone induction.

## References

Azlan A, Zhu L, Fukunaga R. 2024. Female-germline specific protein Sakura interacts with Otu and is crucial for germline stem cell renewal and differentiation and oogenesis. bioRxiv.

Bell LR, Horabin JI, Schedl P, Cline TW. 1991. Positive autoregulation of sex-lethal by alternative splicing maintains the female determined state in Drosophila. Cell 65: 229–239.

Bopp D, Horabin JI, Lersch RA, Cline TW, Schedl P. 1993. Expression of the Sex-lethal gene is controlled at multiple levels during Drosophila oogenesis. Development 118: 797–812.

Chau J, Kulnane LS, Salz HK. 2012. Sex-lethal enables germline stem cell differentiation by down-regulating Nanos protein levels during Drosophila oogenesis. Proceedings of the National Academy of Sciences of the United States of America 109: 9465–9470.

Chen D, McKearin D. 2003a. Dpp signaling silences bam transcription directly to establish asymmetric divisions of germline stem cells. Current biology : CB 13: 1786–1791.

Chen D, McKearin DM. 2003b. A discrete transcriptional silencer in the bam gene determines asymmetric division of the Drosophila germline stem cell. Development 130: 1159–1170.

Chen D, Wang Q, Huang H, Xia L, Jiang X, Kan L, Sun Q, Chen D. 2009. Effete-mediated degradation of Cyclin A is essential for the maintenance of germline stem cells in Drosophila. Development 136: 4133–4142.

Chu HP, Liao Y, Novak JS, Hu Z, Merkin JJ, Shymkiv Y, Braeckman BP, Dorovkov MV, Nguyen A, Clifford PM et al. 2014. Germline quality control: eEF2K stands guard to eliminate defective oocytes. Developmental cell 28: 561–572.

Cox DN, Chao A, Baker J, Chang L, Qiao D, Lin H. 1998. A novel class of evolutionarily conserved genes defined by piwi are essential for stem cell self-renewal. Genes & development 12: 3715–3727.

de Cuevas M, Matunis EL. 2011. The stem cell niche: lessons from the Drosophila testis. Development 138: 2861–2869.

Eliazer S, Shalaby NA, Buszczak M. 2011. Loss of lysine-specific demethylase 1 nonautonomously causes stem cell tumors in the Drosophila ovary. Proceedings of the National Academy of Sciences of the United States of America 108: 7064–7069.

ElMaghraby MF, Tirian L, Senti KA, Meixner K, Brennecke J. 2022. A genetic toolkit for studying transposon control in the Drosophila melanogaster ovary. Genetics 220.

Fukunaga R, Doudna JA. 2009. dsRNA with 5’ overhangs contributes to endogenous and antiviral RNA silencing pathways in plants. The EMBO journal 28: 545–555.

Fukunaga R, Han BW, Hung JH, Xu J, Weng Z, Zamore PD. 2012. Dicer partner proteins tune the length of mature miRNAs in flies and mammals. Cell 151: 533–546.

Gans M, Audit C, Masson M. 1975. Isolation and Characterization of Sex-Linked Female-Sterile Mutants in Drosophila-Melanogaster. Genetics 81: 683–704.

Gateff E, Kurzik-Dumke U, Wismar J, Loffler T, Habtemichael N, Konrad L, Dreschers S, Kaiser S, Protin U. 1996. Drosophila differentiation genes instrumental in tumor suppression. Int J Dev Biol 40: 149–156.

Glenn LE, Searles LL. 2001. Distinct domains mediate the early and late functions of the Drosophila ovarian tumor proteins. Mechanisms of development 102: 181–191.

Gollin SM, King RC. 1981. Studies of Fs(1)1621, a Mutation Producing Ovarian-Tumors in Drosophila-Melanogaster. Developmental Genetics 2: 203–218.

Grmai L, Pozmanter C, Van Doren M. 2022. The Regulation of Germline Sex Determination in Drosophila by Sex lethal. Sex Dev 16: 323–328.

Handler D, Meixner K, Pizka M, Lauss K, Schmied C, Gruber FS, Brennecke J. 2013. The genetic makeup of the Drosophila piRNA pathway. Molecular cell 50: 762–777.

Hayashi Y, Yoshinari Y, Kobayashi S, Niwa R. 2020. The regulation of Drosophila ovarian stem cell niches by signaling crosstalk. Curr Opin Insect Sci 37: 23–29.

Hinnant TD, Merkle JA, Ables ET. 2020. Coordinating Proliferation, Polarity, and Cell Fate in the Drosophila Female Germline. Front Cell Dev Biol 8: 19.

Huang X, Fejes Toth K, Aravin AA. 2017. piRNA Biogenesis in Drosophila melanogaster. Trends Genet 33: 882–894.

Inoue K, Hoshijima K, Sakamoto H, Shimura Y. 1990. Binding of the Drosophila sex-lethal gene product to the alternative splice site of transformer primary transcript. Nature 344: 461–463.

Ji S, Li C, Hu L, Liu K, Mei J, Luo Y, Tao Y, Xia Z, Sun Q, Chen D. 2017. Bam-dependent deubiquitinase complex can disrupt germ-line stem cell maintenance by targeting cyclin A. Proceedings of the National Academy of Sciences of the United States of America 114: 6316–6321.

Ji S, Luo Y, Cai Q, Cao Z, Zhao Y, Mei J, Li C, Xia P, Xie Z, Xia Z et al. 2019. LC Domain-Mediated Coalescence Is Essential for Otu Enzymatic Activity to Extend Drosophila Lifespan. Molecular cell 74: 363–377 e365.

Jin Z, Flynt AS, Lai EC. 2013. Drosophila piwi mutants exhibit germline stem cell tumors that are sustained by elevated Dpp signaling. Current biology : CB 23: 1442–1448.

Jumper J, Evans R, Pritzel A, Green T, Figurnov M, Ronneberger O, Tunyasuvunakool K, Bates R, Zidek A, Potapenko A et al. 2021. Highly accurate protein structure prediction with AlphaFold. Nature 596: 583–589.

Kahney EW, Snedeker JC, Chen X. 2019. Regulation of Drosophila germline stem cells. Current opinion in cell biology 60: 27–35.

Kandasamy SK, Fukunaga R. 2016. Phosphate-binding pocket in Dicer-2 PAZ domain for high-fidelity siRNA production. Proceedings of the National Academy of Sciences of the United States of America 113: 14031–14036.

Kandasamy SK, Zhu L, Fukunaga R. 2017. The C-terminal dsRNA-binding domain of Drosophila Dicer-2 is crucial for efficient and high-fidelity production of siRNA and loading of siRNA to Argonaute2. Rna 23: 1139–1153.

Kang I, Choi Y, Jung S, Lim JY, Lee D, Gupta S, Moon W, Shin C. 2018. Identification of target genes regulated by the Drosophila histone methyltransferase Eggless reveals a role of Decapentaplegic in apoptotic signaling. Scientific reports 8: 7123.

King RC, Riley SF. 1982. Ovarian Pathologies Generated by Various Alleles of the Otu Locus in Drosophila-Melanogaster. Developmental Genetics 3: 69–89.

Kirilly D, Xie T. 2007. The Drosophila ovary: an active stem cell community. Cell research 17: 15–25.

Levin HL, Moran JV. 2011. Dynamic interactions between transposable elements and their hosts. Nature Reviews Genetics 12: 615–627.

Li Y, Minor NT, Park JK, McKearin DM, Maines JZ. 2009. Bam and Bgcn antagonize Nanos-dependent germ-line stem cell maintenance. Proceedings of the National Academy of Sciences of the United States of America 106: 9304–9309.

Li Y, Zhang Q, Carreira-Rosario A, Maines JZ, McKearin DM, Buszczak M. 2013. Mei-p26 cooperates with Bam, Bgcn and Sxl to promote early germline development in the Drosophila ovary. PloS one 8: e58301.

Liao SE, Kandasamy SK, Zhu L, Fukunaga R. 2019. DEAD-box RNA helicase Belle post-transcriptionally promotes gene expression in an ATPase activity-dependent manner. Rna.

Lin H. 1997. The tao of stem cells in the germline. Annu Rev Genet 31: 455–491.

Lin H, Spradling AC. 1993. Germline stem cell division and egg chamber development in transplanted Drosophila germaria. Developmental biology 159: 140–152.

Losick VP, Morris LX, Fox DT, Spradling A. 2011. Drosophila stem cell niches: a decade of discovery suggests a unified view of stem cell regulation. Developmental cell 21: 159–171.

McKearin DM, Spradling AC. 1990. bag-of-marbles: a Drosophila gene required to initiate both male and female gametogenesis. Genes & development 4: 2242–2251.

Mercer M, Dasgupta A, Pawlowski K, Buszczak M. 2025. Bourbon and Mycbp function with Otu to promote Sxl protein expression in the Drosophila female germline. Proceedings of the National Academy of Sciences of the United States of America 122: e2426524122.

Moon S, Cassani M, Lin YA, Wang L, Dou K, Zhang ZZ. 2018. A Robust Transposon-Endogenizing Response from Germline Stem Cells. Developmental cell 47: 660–671 e663.

Ohlstein B, Lavoie CA, Vef O, Gateff E, McKearin DM. 2000. The Drosophila cystoblast differentiation factor, benign gonial cell neoplasm, is related to DExH-box proteins and interacts genetically with bag-of-marbles. Genetics 155: 1809–1819.

Ota R, Kobayashi S. 2020. Myc plays an important role in Drosophila P-M hybrid dysgenesis to eliminate germline cells with genetic damage. Commun Biol 3: 185.

Pauli D, Oliver B, Mahowald AP. 1993. The role of the ovarian tumor locus in Drosophila melanogaster germ line sex determination. Development 119: 123–134.

Pek JW, Lim AK, Kai T. 2009. Drosophila maelstrom ensures proper germline stem cell lineage differentiation by repressing microRNA-7. Developmental cell 17: 417–424.

Penalva LO, Sanchez L. 2003. RNA binding protein sex-lethal (Sxl) and control of Drosophila sex determination and dosage compensation. Microbiology and molecular biology reviews : MMBR 67: 343–359, table of contents.

Rodesch C, Pettus J, Nagoshi RN. 1997. The Drosophila ovarian tumor gene is required for the organization of actin filaments during multiple stages in oogenesis. Developmental biology 190: 153–164.

Rubin T, Huynh JR. 2015. Mosaic Analysis in the Drosophila melanogaster Ovary. Methods in molecular biology 1328: 29–55.

Salz HK, Erickson JW. 2010. Sex determination in Drosophila: The view from the top. Fly 4: 60–70.

Smith PA, King RC. 1966. Studies on Fused a Mutant Gene Producing Ovarian Tumors in Drosophila Melanogaster. Genetics 54: 363-&.

Steinhauer WR, Kalfayan LJ. 1992. A Specific Ovarian Tumor Protein Isoform Is Required for Efficient Differentiation of Germ-Cells in Drosophila Oogenesis. Genes & development 6: 233–243.

Storto PD, King RC. 1988. Multiplicity of functions for the otu gene products during Drosophila oogenesis. Dev Genet 9: 91–120.

Taira T, Maeda J, Onishi T, Kitaura H, Yoshida S, Kato H, Ikeda M, Tamai K, Iguchi-Ariga SM, Ariga H. 1998. AMY-1, a novel C-MYC binding protein that stimulates transcription activity of C-MYC. Genes Cells 3: 549–565.

Taira Y, Zhu L, Fukunaga R. 2025. RNA-binding protein Miso/CG44249 is crucial for minor splicing during oogenesis in Drosophila. Rna 31: 822–835.

Valcarcel J, Singh R, Zamore PD, Green MR. 1993. The protein Sex-lethal antagonizes the splicing factor U2AF to regulate alternative splicing of transformer pre-mRNA. Nature 362: 171–175.

Wang Z, Lin H. 2004. Nanos maintains germline stem cell self-renewal by preventing differentiation. Science 303: 2016–2019.

Xia L, Jia S, Huang S, Wang H, Zhu Y, Mu Y, Kan L, Zheng W, Wu D, Li X et al. 2010. The Fused/Smurf complex controls the fate of Drosophila germline stem cells by generating a gradient BMP response. Cell 143: 978–990.

Xie T, Spradling AC. 1998. decapentaplegic is essential for the maintenance and division of germline stem cells in the Drosophila ovary. Cell 94: 251–260.

Xu T, Rubin GM. 1993. Analysis of genetic mosaics in developing and adult Drosophila tissues. Development 117: 1223–1237.

Yamashiro H, Siomi MC. 2018. PIWI-Interacting RNA in Drosophila: Biogenesis, Transposon Regulation, and Beyond. Chem Rev 118: 4404–4421.

Yang F, Quan Z, Huang H, He M, Liu X, Cai T, Xi R. 2019. Ovaries absent links dLsd1 to HP1a for local H3K4 demethylation required for heterochromatic gene silencing. eLife 8.

Yang L, Chen D, Duan R, Xia L, Wang J, Qurashi A, Jin P, Chen D. 2007. Argonaute 1 regulates the fate of germline stem cells in Drosophila. Development 134: 4265–4272.

Yang Z, Zeng X, Zhao Y, Chen R. 2023. AlphaFold2 and its applications in the fields of biology and medicine. Signal Transduct Target Ther 8: 115.

Zhu L, Fukunaga R. 2021. RNA-binding protein Maca is crucial for gigantic male fertility factor gene expression, spermatogenesis, and male fertility, in Drosophila. PLoS genetics 17: e1009655.

Zhu L, Kandasamy SK, Fukunaga R. 2018a. Dicer partner protein tunes the length of miRNAs using base-mismatch in the pre-miRNA stem. Nucleic acids research.

Zhu L, Kandasamy SK, Liao SE, Fukunaga R. 2018b. LOTUS domain protein MARF1 binds CCR4-NOT deadenylase complex to post-transcriptionally regulate gene expression in oocytes. Nature communications 9: 4031.

Zhu L, Liao SE, Ai Y, Fukunaga R. 2019a. RNA methyltransferase BCDIN3D is crucial for female fertility and miRNA and mRNA profiles in Drosophila ovaries. PloS one 14: e0217603.

Zhu L, Liao SE, Fukunaga R. 2019b. Drosophila Regnase-1 RNase is required for mRNA and miRNA profile remodelling during larva-to-adult metamorphosis. RNA biology 16: 1386–1400.

